# Gene swamping alters evolution during range expansions in the protist *Tetrahymena thermophila*

**DOI:** 10.1101/863340

**Authors:** Felix Moerman, Emanuel A. Fronhofer, Andreas Wagner, Florian Altermatt

## Abstract

At species’ range edges, individuals often face novel environmental conditions that may limit range expansion until populations adapt. The potential to adapt depends on genetic variation upon which selection can act. However, populations at species’ range edges are often genetically depauperated. One mechanism to increase genetic variation is to reshuffle existing variation through sex. During range expansions, sex can, however, act as a double-edged sword. The gene swamping hypothesis predicts that for populations expanding along an abiotic gradient, sex can hinder adaptation if asymmetric dispersal leads to numerous maladapted dispersers from the range core swamping the range edge. In this study, we experimentally tested the gene swamping hypothesis by performing replicated range expansions in landscapes with or without an abiotic pH-gradient, using the ciliate *Tetrahymena thermophila*, while simultaneously manipulating the occurrence of gene flow and sex. We show that sex accelerated evolution of local adaptation in the absence of gene flow, but hindered it in the presence of gene flow. The effect of sex, however, was independent of the pH-gradient, indicating that not only abiotic gradients but also the biotic gradient in population density leads to gene swamping. Overall, our results show that gene swamping can affect adaptation in life-history strategies.

## Introduction

Individuals living at the edge of a species’ range face different conditions compared to those in the core region. Selection pressures differ, and often the individuals at the edge represent only a small subset of a species’ genetic variation [1]. The potential of a population to spread depends on the capacity to disperse and the ability to grow in the local abiotic environment [2]. Consequently, when populations expand their range, they experience strong selection due to the range expansion itself, and are also affected by concurrently changing environmental conditions.

During range expansions, populations can undergo rapid evolution, as demonstrated by recent comparative and experimental work [1], showing evolution of increased dispersal [3, 4, 5, 6], r-selected life-history strategies [7, 8], and adaptation to abiotic conditions [9, 10]. Expanding into previously uninhabited space allows populations to escape intraspecific competition. Consequently, evolving in response to multiple selective pressures can potentially lead to substantial benefits, despite the challenges involved [11, 8].

A major modulator of evolution is sex. Sex allows populations to reshuffle existing genetic variation [12, 13, 14, 15]. Theoretical work suggests that sex would typically lead to offspring with lower fitness, by breaking up advantageous allele combinations (recombination load), and hence an advantage for asexual reproduction [16]. However, populations during range expansion experience strong stochasticity due to repeated founder events, leading to maladaptive mutations becoming fixed and surfing along at the range edge (expansion load) [17, 18]. Sex can strongly reduce these negative effects of expansion load, thus making it advantageous [19, 18, 20].

If populations face strong abiotic stressors or heterogeneous environments, sex may also facilitate adaptation [21, 22, 23]. Given that experimental work found stronger benefits of sex if genetic variation is sufficiently high [24], we expect that sex is only favoured at the range edge when genetic variation is bolstered through gene flow from the high diversity core, because populations at a range edge are genetically depauperated due to repeated founder events [1, 25]. However, theory on gene swamping predicts the opposite [26, 27, 28]. As individuals bolstering the gene pool will be maladapted to the abiotic conditions at the range edge, sex may hinder adaptation when there is too much gene flow from the range core to the range edge [26, 27, 28, 29, 30]. Under such conditions, reproducing sexually would swamp the gene pool at the range edge with maladapted genes. This could prevent the population from adapting to the abiotic environment at the range edge, and hence slow down and even halt range expansion, leading to stable range borders [27, 28]. In contrast, when drift strongly reduces adaptive variation, gene flow may positively affect adaptation by counteracting the effects of drift [29, 30]. Despite extensive theory on gene swamping, surprisingly little empirical and experimental work exists [31, 32, 33, 34].

Here, we experimentally tested the gene swamping hypothesis using the ciliate *Tetrahymena thermophila*. We assessed how reproduction (asexual or sexual) and gene flow (i.e., dispersal from the range core to the range edge) altered evolutionary adaptation during range expansions in landscapes with or without a gradient in pH. We found a distinct signal of gene swamping, where sex facilitated or hindered adaptation depending on the presence or absence of gene flow.

## Material and methods

### Study organism

*Tetrahymena thermophila* is a freshwater ciliate commonly used in ecological and evolutionary experiments [35, 36, 37, 38, 39]. We used four phenotypically divergent [40] clonal strains of *T. thermophila* obtained from the Tetrahymena Stock Center: strain B2086.2 (Research Resource Identifier TSC SD00709), strain CU427.4 (TSC SD00715), strain CU428.2 (TSC SD00178) and strain SB3539 (TSC SD00660).

### Experiment

#### Microcosms

We performed all evolution experiments and all bioassays in a 20 °C climate-controlled room. Following an established method [4], we experimentally emulated an expanding range front with two-patch landscapes, which consisted of two 25 mL Sarstedt tubes connected by an 8 cm long silicone tube (inner diameter 4 mm). See also Supplementary Material figure S1.

We prepared 40 two-patch landscapes, and filled patches of each landscape with 15 mL modified Neff-medium [41]. We complemented the medium for experimental evolution and bioassays with 10 μgmL^−1^ Fungin and 100 μgmL^−1^ Ampicillin to prevent bacterial and fungal contamination. We then inoculated one patch of each two-patch landscape with 200 μL of ancestor culture (50 μL from each of the four ancestral strains). This allowed adaptation through clonal selection and de novo mutation [42] in populations designated for asexual reproduction, as well as recombination [43] in populations designated for sexual reproduction.

#### Treatment groups

We designed a full-factorial experiment that tested the effect of 1) abiotic conditions, with two treatment levels (“Uniform”: pH always 6.5, “Gradient”: pH starts at 6.5 and then gradually decreases), 2) reproduction, with two treatment levels (“Asexual”: pure asexual reproduction, “Sexual”: asexual and sexual reproduction) and 3) gene flow, with two treatment levels (“Absent”: no gene flow; “Present”: gene flow from the range core to range edge). We evolved five replicate populations per treatment, for a total of 40 evolving populations.

#### Experimental evolution

We performed a range expansion experiment that lasted ten weeks, in which we repeated the same procedure cycle every 14 days. This cycle consisted of three dispersal events (on days 1, 3 and 5). These events were followed by a gene flow and sexual reproduction event or the appropriate controls depending on the treatment groups (on day 8), and subsequently an additional two dispersal events (on days 10 and 12).

We initiated dispersal by opening the clamps in the two-patch landscapes for one hour, which allowed cells to disperse from their original (home) patch to the target patch. After dispersal, we prepared 40 new two-patch landscapes. If population density was measurable (≥1 cell observed during video analysis, see below) in the target patch, we transferred the content of the target patch to a new two-patch landscape. If no measurable dispersal occurred, we transferred the content of the home patch to the new two-patch landscape.

In treatment groups designated for gene flow to occur, we emulated long-distance gene flow (from the range core to the edge, following theoretical predictions [27, 28]), by transferring 1.5 mL of culture from the core population to the range front.

To control reproduction, we transferred all populations to a starvation medium, because *T. thermophila* only mates when starved [43]. We incubated the starvation cultures on a shaker rotating at 120 rpm. After 36 hours, we placed the populations designated for sexual reproduction off the shaker, but kept populations designated for asexual reproduction on the shaker, because the shaking movement prevents cells from mating. We left cells to mate overnight, after which we transferred populations to new two-patch landscapes. For a more extensive technical description, see Supplementary Material section S1.2.

#### Common garden

After experimental evolution, we sampled 100 μL of culture from all surviving populations, and transferred this sample to 25 mL Sarstedt tubes containing 15 mL Neff-medium at pH 6.5. We maintained these populations in the common garden for 72 hours before starting bioassays, to reduce epigenetic and trans-generational effects.

#### Bioassays

We quantified the population growth rate of ancestral and evolved populations, after common garden cultivation, at eight different pH values (pH 6.5, 6.0, 5.5, 5.0, 4.5, 4.0, 3.5 and 3.0). Specifically, we prepared for every population Sarstedt tubes containing Neff-medium whose pH we had adjusted to the desired value using 1 M HCL, and inoculated this medium with 100 μL of culture from the evolved or ancestral populations. We grew the resulting cultures for 12 days, sampling populations twice on the first two days, and once per day on all subsequent days. Every two days, we replaced 1 mL of culture with fresh medium to prevent population decline.

#### Sampling and video analysis

We measured population density and cell characteristics (morphology and movement) using an established method [36, 44]. We sampled 200 μL of culture from every population, and diluted samples 10—100 fold in Neff-medium to ensure densities were similar, as excessive density prevents accurate video analysis. We then took 10 s videos (250 frames, 25 fps) using a Leica M165FC stereomicroscope and top-mounted Hamamatsu Orca Flash 4.0 camera. We analyzed videos using the BEMOVI R-package [44] (parameters in Supplementary Material section S2).

### Beverton-Holt model fitting

To analyze local adaptation, we assessed growth rates by fitting a continuous-time version of the Beverton-Holt model [45], as this model is well-suited for microcosm data and facilitates biological interpretation of parameters [46, 47]. The Beverton-Holt model is given by the equation:

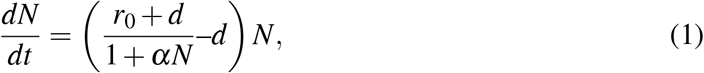

where the intraspecific competitive ability (*α*) is equal to

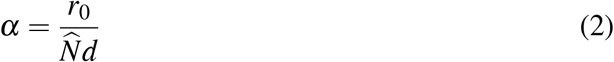

and *r*_0_ is the intrinsic rate of increase, *N* the population size, *α* the intraspecific competitive ability, 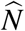 the equilibrium population density and *d* the death rate of the population. We estimated the parameters using a Bayesian approach adapted from Rosenbaum et al. [48]. For model code see https://zenodo.org/record/2658131

### Statistical analysis

All statistical analyses were performed with the R language for statistical computing, version 3.5.1. We calculated local adaptation by assessing changes in the intrinsic rate of increase *r*_0_ of evolved populations under the pH conditions they experienced during evolution, compared to the ancestor under the same pH conditions. This was done by dividing the *r*_0_ estimates of evolved populations by the mean *r*_0_ of the mixed ancestral populations (populations with the initial ancestral genotype mixture), and by subsequently calculating the logarithm (base 2) of this ratio (log-ratio response).

Next, we created linear models assessing the effect of reproduction, gene flow and abiotic conditions (explanatory variables) on range expansion distance (number of successful dispersal events) and local adaptation respectively. We additionally created a linear mixed model (‘nlme’-package, version 3.1-137) to assess how population density during range expansion was influenced by the three treatments: reproduction, gene flow, abiotic conditions, as well as the covariate range expansion distance (the number of successful dispersal events). We included population ID as a random effect. We subsequently compared all possible models for these three response variables using the dredge function (‘MuMin’-package, version 1.43.6) to select the model with lowest AICc (Akaike Information Criterion, corrected for small sample size [49]) score for local adaptation and range expansion distance, and lowest BIC (Bayesian information criterion [50]) for population density. We report relative importance and model output. See Supplementary Material section S4 for additional analyses on population survival and cell movement and morphology.

## Results

Population densities (figure 1 panels A, B, E and F; Table 1) showed strong temporal variation in all replicates. Mean density decreased marginally for populations expanding into uniform abiotic conditions (*χ*^2^_1,746_=4.526, p=0.034), whereas population density of populations expanding into a gradient decreased strongly (*χ*^2^_1,746_=108.258, p<0.0001). Additionally, we observed that populations faced with a gradient showed significantly slower range expansion (figure 1 panels C, D, G and H; F_1,31_=141.4, p<0.0001; table 2), and were more prone to go extinct, in the absence of gene flow (see Supplementary material section S4.6).

**Table 1:** Type III ANOVA table of the best model for population density during range expansion according to BIC model comparison.

| Model and explanatory variables | Degrees of freedom | $\chi^2$-value | Pr ($>\chi^2$) |
| --- | --- | --- | --- |
| Abiotic conditions | 1 | 0.044 | 0.833 |
| Range expansion distance | 1 | 4.526 | 0.034 |
| Abiotic conditions × position | 1 | 108.258 | <.0001 |

**Table 2:** Type III ANOVA table of the best model for local adaptation (evolution of intrinsic rate of increase *r*_0_) and range expansion distance (total number of successful dispersal events) during range expansion according to AICc model comparison.

| Model and explanatory variables | Degrees of freedom | F-value | Pr (>F) |
| --- | --- | --- | --- |
| **Local adaptation** |  |  |  |
| Reproduction | 1 | 3.96 | 0.056 |
| Gene flow | 1 | 5.55 | 0.025 |
| Abiotic conditions | 1 | 122.58 | <0.0001 |
| Reproduction×Gene flow | 1 | 10.67 | 0.003 |
| Residuals | 29 |  |  |
| **Range expansion distance** |  |  |  |
| Abiotic conditions | 1 | 141.4 | <0.0001 |
| Residuals | 31 |  |  |

**Figure 1:**
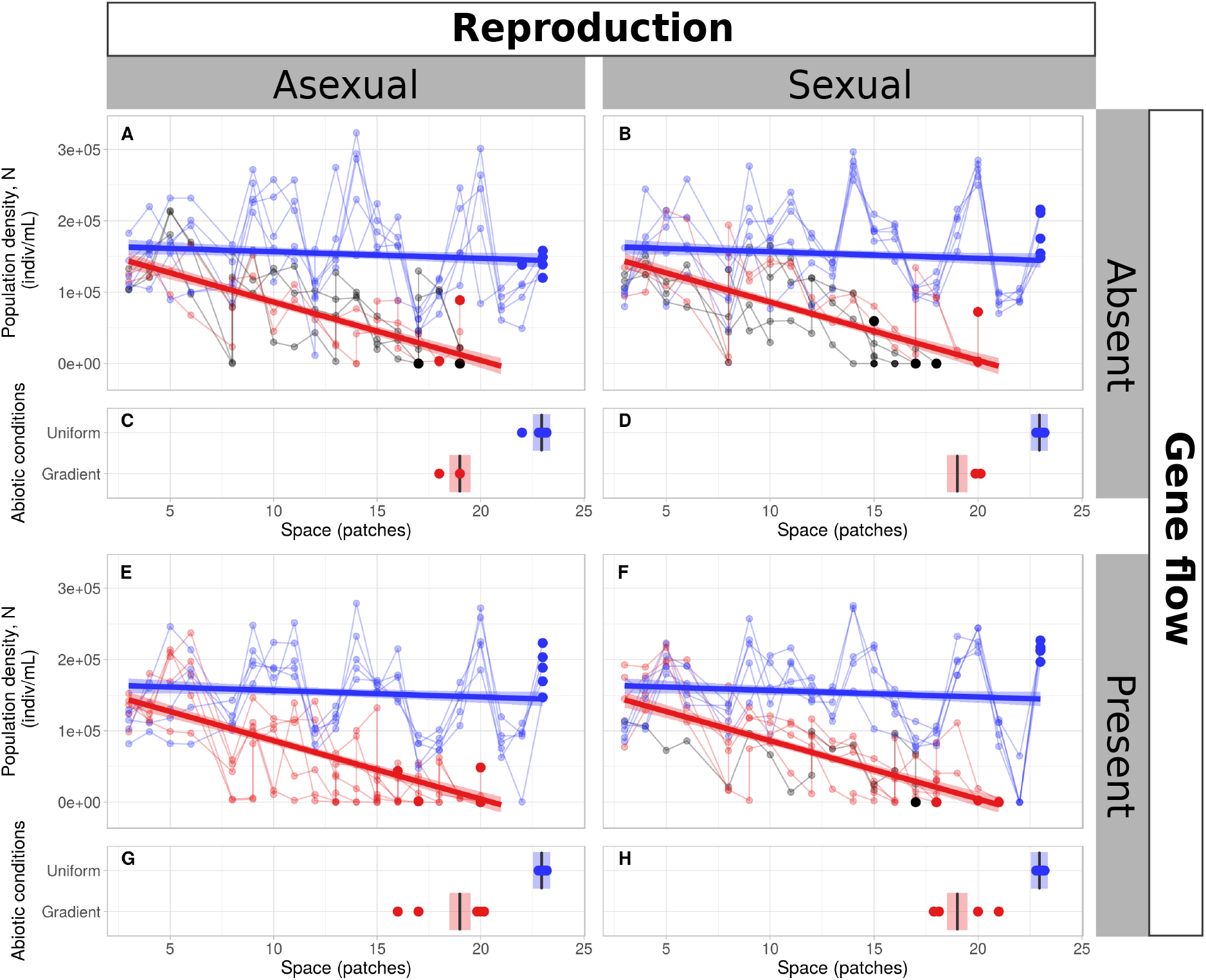
Population dynamics for the different treatment groups over the course of the range expansion dynamics. Faint blue lines and dots represent data for the populations expanding into uniform abiotic conditions. Faint red lines and dots show the data for populations expanding into a gradient (only given for the populations that survived until the bioassays). Faint black lines and dots show data for populations expanding into a gradient, but went extinct before the start of the bioassays. The larger and brighter coloured dots represent the population densities measured at the last timepoint. Thick lines and shaded areas show the mean model predictions and 95 %-confidence intervals respectively respectively, for the best model (according to BIC/WAIC comparisons through the dredge function) on population densities/range expansion distances of surviving populations expanding into a gradient (red) or uniform abiotic conditions (blue). The large panels (A, B, E and F) show population densities as a function of distance dispersed during the range expansion experiment. The small plots (C, D, G, H) show the data and model predictions on total distance expanded by the end of the range expansion experiment of the surviving populations.

Local adaptation (evolution of intrinsic rate of increase *r*_0_; figure 2; table 2) increased only slightly for population expanding into uniform abiotic conditions, whereas populations that expanded into a gradient greatly increased local adaptation (F_1,29_=128.58, p<0.0001). Although reproduction (F_1,29_=3.96, p=0.056) and gene flow (F_1,29_=5.55, p=0.025) individually slightly increased local adaptation, their interaction strongly decreased local adaptation (F_1,29_=10.67, p=0.003), with populations evolving lower intrinsic rates of increase either when reproduction was sexual and gene flow present, or with asexual reproduction but gene flow absent.

**Figure 2:**
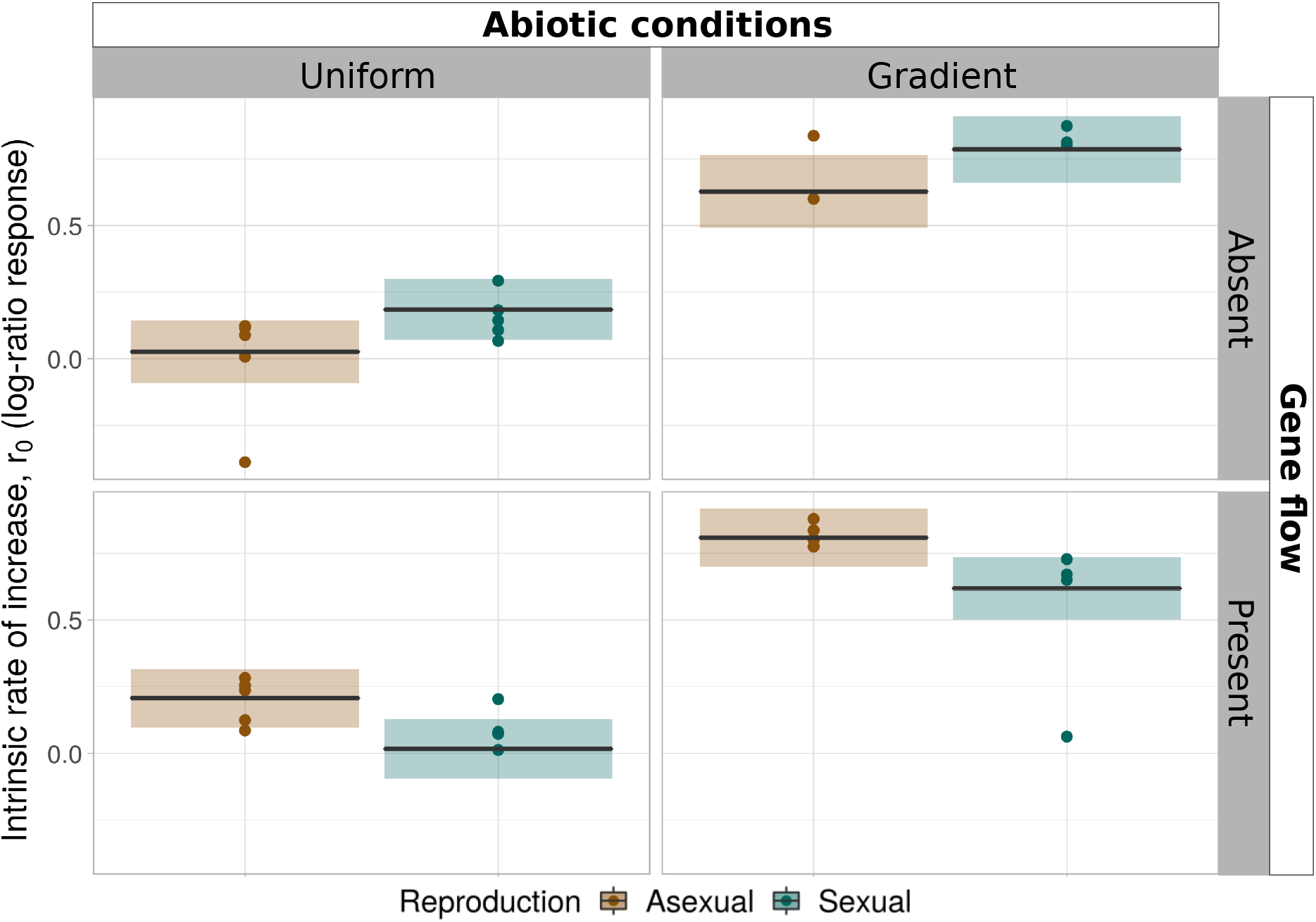
Local adaptation, measured as the evolution of intrinsic rate of increase *r*_0_ in the abiotic conditions experienced during range expansion (uniform or gradient), compared to the ancestor population. The y-axis shows the change in *r*_0_ compared to the ancestor (log-ratio response). Dots represent individual data points, black lines and shaded areas show the model predictions of the best model (mean and 95 %-confidence interval). Brown colours denote populations in treatment groups with asexual reproduction, green colours denote populations in treatment groups with sexual reproduction.

## Discussion

We experimentally assessed the gene swamping hypothesis by performing replicated range expansions using the protist *Tetrahymena thermophila*. We experimentally manipulated abiotic conditions (uniform versus gradient), reproduction (asexual versus sexual) and gene flow (absent versus present). We demonstrated how sex interacts with gene flow, affecting local adaptation of organisms at the range edge (figure 2; table 2).

Populations undergoing range expansions face multiple selective pressures [1], and hence face a strong pressure to adapt. Theoretical predictions suggest that sex can be advantageous or disadvantageous during range expansion, depending on the context. Theory on gene swamping predicts that sex hinders adaptation during range expansions when populations undergo strong asymmetrical dispersal from a range core to a range edge [26, 27, 28]. We showed here that sex and gene flow interact during range expansions, modulating local adaptation. Despite having only four distinct events of sexual reproduction in otherwise asexually reproducing populations, we found that sex facilitated adaptation in the absence of gene flow. However, this effect was reversed when gene flow was present, and swamped the edge population with maladapted individuals. Surprisingly, while the gene swamping hypothesis predicts this pattern exclusively in the presence of abiotic gradients [26, 27, 28], we observed similar effects of gene swamping in the presence and absence of an abiotic gradient. We argue that gene swamping in the absence of an abiotic gradient could stem from evolving life-history strategies during range expansions. Range expanding populations are thought to exhibit a gradient of decreased density towards the range front, which translates to decreased competition and selection for fast reproduction [51]. Hence, gene swamping may imply that individuals maladapted in life-history strategy interbreed with the population at the range edge. Consequently gene swamping affects adaptation during range expansions even without an abiotic gradient, leading to analogous changes in adaptation as for range expansions into abiotic gradients.

Although we show that gene swamping affects adaptation during range expansions, we could not detect effects of gene swamping on range expansion rates as described by theory, despite population growth rate being a driving force behind expansion rate [2, 52]. Such a could result from our experimental setup, where we used discrete landscapes connected through repeated dispersal events, rather than continuous dispersal. This setup may be insufficiently sensitive to detect signals in expansion rate. Alternatively, this setup may lead to pushed rather than pulled waves (see Pachepsky and Levine [53]) which changes predictions. Under pulled waves, dispersers from the low-density range front drive further range expansion. In contrast, further spread in pulled waves will only be possible after the population at the front has grown sufficiently large. Although it is possible that the abiotic gradient leads to a pushed wave, for example by reducing survival during the dispersal stage, determining this with absolute certainty would require extensive dispersal measurements at a temporal resolution that we lack in the current experiment. Testing the interaction between pushed/pulled waves and gene swamping would, however, be interesting, as pushed waves might be less susceptible to gene swamping, because the population density gradient from the range core to the range edge is less steep compared to pulled waves.

## Acknowledgements

We thank Samuel Hürlemann, Silvana Käser and Sarah Bratschi for help with laboratory work, and Lynn Govaert, Claire Jacquet and the two reviewers for their helpful comments. Funding is from the University of Zurich URPP Evolution in Action and the Swiss National Science Foundation, Grant No PP00P3 179089. This is publication ISEM-YYYY-XXX of the Institut des Sciences de l’Evolution – Montpellier. We would also like to acknowledge support by Swiss National Science Foundation grant 31003A 172887 and European Research Council Advanced Grant No. 739874.

## Author contributions

FM, EAF, AW and FA designed the experiment. FM performed experimental work and statistical analyses. FM, FA, AW and EAF interpreted the results. FM and FA wrote the first version of the manuscript and all authors commented on the final version.

## Data accessibility statement

Data is available from the Dryad Digital Repository (DOI: 10.5061/dryad.6wwpzgmtk) and analysis scripts on Github (DOI: 10.5281/zenodo.3560982).

