## Supplementary Material for "Gene swamping alters evolution during range expansions in the protist *Tetrahymena thermophila*"

### S1 Experimental design

#### S1.1 Treatment groups

Table S1: Overview showing the treatments, their treatment levels (column values) and the replication number included in the experimental design.

| | Abiotic conditions | Reproduction | Gene flow | Replicates |
|---|---|---|---|---|
| **Treatment combination 1** | “Uniform” | “Asexual” | “Absent” | 5 |
| **Treatment combination 2** | “Uniform” | “Asexual” | “Present” | 5 |
| **Treatment combination 3** | “Uniform” | “Sexual” | “Absent” | 5 |
| **Treatment combination 4** | “Uniform” | “Sexual” | “Present” | 5 |
| **Treatment combination 5** | “Gradient” | “Asexual” | “Absent” | 5 |
| **Treatment combination 6** | “Gradient” | “Asexual” | “Present” | 5 |
| **Treatment combination 7** | “Gradient” | “Sexual” | “Absent” | 5 |
| **Treatment combination 8** | “Gradient” | “Sexual” | “Present” | 5 |

### S1.2 Extended methodology experimental evolution

Figure S1 shows a graphic representation of the design used for the experimental range expansion. Panel A shows the time course of experimental evolution on the vertical axis. The horizontal axis indicates the distance along which populations expanded during the experiment. The leftmost section between the vertical axis and the solid vertical line shows the four ancestral populations (depicted as 4 separate tubes/microcosms). We kept these four ancestral populations at a neutral pH (6.5) conditions as slowly dividing cultures over the entire course of the experiment. Because these conditions were identical to the conditions in which the ancestral populations had been kept before the start of the experiment, we do not expect directed evolutionary change in these ancestral populations. Note, however, that although we maintain these four ancestral clones separately, we initiated the actual range expansion experiment with a mixture of the four ancestors. Keeping the four ancestral populations separately allowed us to mix the four ancestors at the exact same proportions as used at the start of the experiment to emulate gene flow during the experiment. On the right side of the vertical solid line, the figure shows the expanding range front. Because we only tracked the very front of the range expansion using our two-patch landscape, we indicate the front through a pair of brightly coloured and connected culture vessels, which represent two-patch landscapes. During the range expansion experiment, whenever populations successfully dispersed, the range front moves to the right, depicted here by the rightward shift of a brightly coloured culture vessel pair. By only tracking the very front of the range expansions, earlier intermediate patches were no longer maintained. We depict these no longer maintained locations as faintly coloured culture vessels. These faint culture vessels thus represent positions where the range expansion front was located in previous timepoints of the experiment. They are merely shown to help visualize the spatial aspect of the range expansion.

The timeline is divided into 7 distinct horizontal sections (demarcated by the numbers 1-7 in square brackets). The first section (labeled “Start experiment”) shows how we initiated the experiment, namely by inoculating the first tube of a two-patch landscape with 50 µL culture volume from each of the four ancestral populations. The second section (Cycle 1) shows the steps involved in each of the five 14-day cycles. The steps in each cycle consisted of three dis-

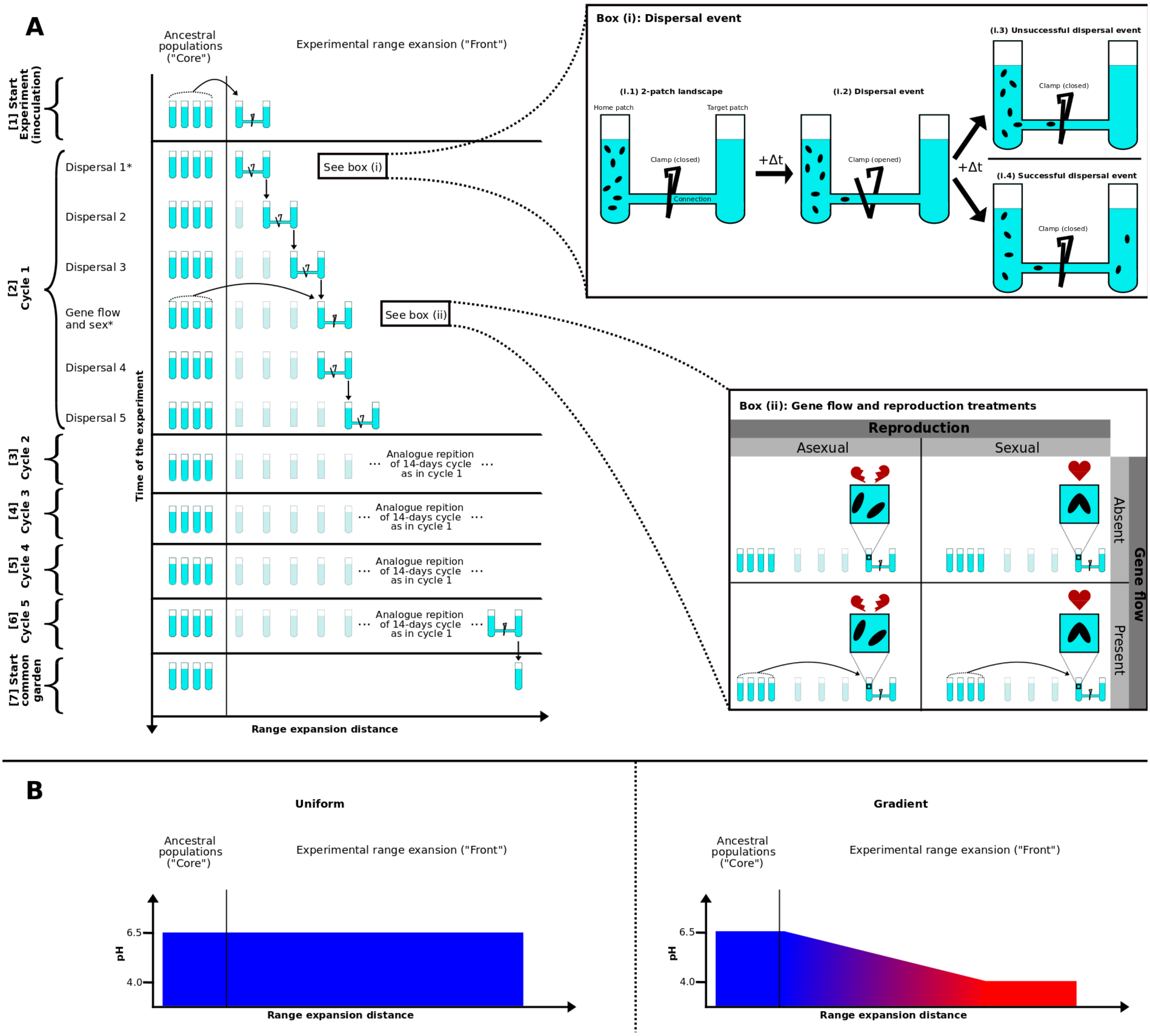


Figure S1: Schematic representation of the experimental range expansion. Panel A shows the timeline of the experiment, with the y-axis depicting time, and the x-axis the distance populations expanded during range expansion. Box (i) shows in more detail the experimental approach involved in a dispersal step, and box (ii) the treatments related to gene flow and reproduction. Panel B shows the abiotic conditions experienced during range expansion into a gradient or uniform environment.

persal events, followed by gene flow and sexual reproduction (in the relevant treatment groups), and two further dispersal events.Whereas a typical 14-day cycle is shown in Cycle 1, we made two exceptions to these steps, which are marked in figure S1 by an asterisk. In Cycle 1, we skipped the dispersal 1 step, as this dispersal step would have taken place on the same day as the initiation of the experiment. Population densities at this time were too low for measurable dispersal to take place. Consequently, we allowed the populations to grow for two days after the initiation, before initiating the first dispersal event (dispersal 2). The second exception regards to Cycle 5, in which we did not start another gene flow and sexual reproduction event, as this would have taken place too close to the end of the experiment. Sections 3-6 (Cycle 2-5) of the panel depict how we repeated the initial 14-day cycle an additional four times during the experiment.

The dispersal and gene flow/sexual reproduction events are shown in more detail in boxes (i) and (ii) of figure S1. Before a dispersal event (box i.1), all cells (depicted in the box as black ovals) are confined to the first tube (the home patch) of a two-patch landscapes, because plastic clamps keep the connection between the two patches closed. At the start of the dispersal event (box i.2), we opened this clamp and allowed cells to actively swim between the home patch and the target patch. We left the clamp open for one hour. After closure of the clamp, we measured population densities in the home patch and target patch, as described in the main text (Section "Sampling and video analysis"). If no measurable number of cells had dispersed to the target patch of the two-patch landscape (box i.3), we assumed that dispersal was unsuccessful, and we transferred the content of the home patch to a new two-patch landscape. When some cells had successfully dispersed (box i.4), we transferred the content of the target patch to a new two-patch landscape. At the end of the range expansions experiment, the total number of successful dispersal events was used to calculate the total range expansion distance as the sum of the number successful dispersal events.

Box (ii) of figure S1 shows the four treatments related to gene flow and sex. To permit gene flow (bottom panels in box (ii)), we removed 1.5 mL of culture from the population, and replaced it with 1.5 mL of the initial (ancestral) culture, containing the four ancestral strains. We prepared this replacement culture by mixing culture of the four ancestral clones so that population densities of each of the four ancestors was the same as the population densities used at the start of the experiment, so that the replacement mixture mimicked the starting conditions of the experiment. For populations in which we did not allow gene flow (top panels of box (ii)), we simply did not perform this replacement.

After the gene flow step, we transferred all populations to starvation medium, because starvation triggers mating in *T. thermophila*. Specifically, we first transferred all populations to 15 mL Falcon tubes, and centrifuged them (5 min, 1100 G, 4 °C in order to pellet the cells, and discarded the supernatant. We subsequently resuspended cells in 10 mM Tris-HCl, with pH adjusted to 7.5. We then repeated the same centrifugation step, and again resuspended cells in 10 mM Tris-HCl with pH adjusted to 7.5. After the second centrifugation, we transferred the populations to autoclaved 100 mL Erlenmeyer bottles, and placed them for approximately 36 hours on a shaker rotating at 120 rpm to starve the cells. On the next day, we allowed sexual reproduction to occur in those populations designated for sexual reproduction (right panels in box (ii) of figure S1), by simply stopping shaking, which allows cells to conjugate (blue squares in box (ii) of figure S1, where cells join). The populations designated for asexual reproduction (left panels in box (ii) of figure S1) remained on the shaker, whose movement prevented cells from successfully conjugating (blue squares in which cells remain separate). We had tested this protocol during pilot experiments). In shaken populations, no cell conjugation occurred, whereas in shaken populations, almost all cells conjugate (data not shown).

After having left cells to mate overnight, we transferred all populations to new 15 mL Falcon tubes. We centrifuged the populations (5 min, 1100 G, 4 °C), removed the supernatant and resuspended cells in Neff medium with its pH adjusted to a specific value using 1 M HCl - either pH 6.5 for populations expanding into uniform abiotic conditions, or increasingly low pH for populations expanding into a gradient. We then transferred these cultures to fresh two-patch landscapes, and allowed them to rest for approximately one hour, before initiating the next dispersal event.

After the end of Cycle 5, we cultivated the populations that had not become extinct during the experiment (34 out of 40 initial population) in a common garden environment ("Start common garden" in panel A of figure S1). We classified a population as extinct if we did not find any living cells in either the home patch or the target patch of the 2-patch landscape. From every surviving population, we extracted culture from either the target patch of the two-patch landscape if the last dispersal event had been successful, or the home patch if dispersal was unsuccessful. We prepared common garden populations by adding 15 mL of Neff-medium to 25 mL Sarstedt tubes, and adding 100 µL of culture from the appropriate population. Cultures remained in the common garden for approximately 72 hours, before starting the bioassays.

Panel B of figure S1 shows the abiotic conditions associated with the uniform and gradient treatment levels. For the uniform treatment level, a population experienced neutral pH of 6.5 (depicted as blue colour) everywhere along its expanding range. For the Gradient treatment level, the environment became increasingly harsher (lower pH; red colour) during range expansion, until a minimum of pH 4.0 was reached. After reaching this minimum, the pH remained constant for the remainder of the range expansion. We gradually lowered the pH in the Gradient treatment, by adjusting the pH of the target patch in steps of 0.5 every 2-3 successful dispersal events, so that if dispersal was always successful, the pH would decrease by 0.5 every week. We kept the ancestral populations as slow-dividing cultures under neutral pH conditions (pH 6.5) over the entire course of the experiment.

#### S1.3 Experimental handling

Table S2: Overview of the experimental handling operations performed on the different days during the range expansion experiment.

| **Day of experiment** | **Handling** |
|---|---|
| 1 | Start experiment |
| 2 | |
| 3 | Dispersal |
| 4 | |
| 5 | Dispersal |
| 6 | |
| 7 | |
| 8 | Gene Flow + Starvation |

| | |
|---|---|
| 9 | Sex |
| 10 | Dispersal |
| 11 | |
| 12 | Dispersal |
| 13 | |
| 14 | |
| 15 | Dispersal |
| 16 | |
| 17 | Dispersal |
| 18 | |
| 19 | Dispersal |
| 20 | |
| 21 | |
| 22 | Gene Flow + Starvation |
| 23 | Sex |
| 24 | Dispersal |
| 25 | |
| 26 | Dispersal |
| 27 | |
| 28 | |
| 29 | Dispersal |
| 30 | |
| 31 | Dispersal |
| 32 | |
| 33 | Dispersal |
| 34 | |
| 35 | |
| 36 | Gene Flow + Starvation |
| 37 | Sex |

Dispersal

Dispersal

Dispersal

Dispersal

Dispersal

Gene Flow + Starvation

Sex

Dispersal

Dispersal

Dispersal

Dispersal

Dispersal

Dispersal

Dispersal

| | |
|---|---|
| 67 | |
| 68 | Transfer to common garden setting |

### S2 Video analysis script

Below follows the script used for video analysis, including all the used parameters.

```
########################################################################
# R script for analysing video files with BEMOVI (www.bemovi.info)
rm(list=ls())
# load package
library(devtools)
install_github("efronhofer/bemovi", ref="experimental")
library(bemovi)

########################################################################
# VIDEO PARAMETERS

# video frame rate (in frames per second)
fps <- 25
# length of video (in frames)
total_frames <- 500

# measured volume (in microliter)
measured_volume <- 34.4 # for Leica M205 C with 1.6 fold magnification,
sample height 0.5 mm and Hamamatsu Orca Flash 4

# size of a pixel (in micrometer)
pixel_to_scale <- 4.05 # for Leica M205 C with 1.6 fold magnification,
sample height 0.5 mm and Hamamatsu Orca Flash 4
```

```
# specify video file format (one of "avi","cxd","mov","tiff")
# bemovi only works with avi and cxd. other formats are reformated to avi below
video.format <- "cxd"

# setup
difference.lag <- 10
thresholds <- c(10,255) # don't change the second value
#thresholds <- c(50,255)

######################################################################
# FILTERING PARAMETERS
# min and max size: area in pixels
particle_min_size <- 5
particle_max_size <- 1000

# number of adjacent frames to be considered for linking particles
trajectory_link_range <- 3
# maximum distance a particle can move between two frames
trajectory_displacement <- 16

# these values are in the units defined by the parameters above: fps (seconds),
#measured_volume (microliters) and pixel_to_scale (micometers)
filter_min_net_disp <- 25
filter_min_duration <- 1
filter_detection_freq <- 0.1
filter_median_step_length <- 3

######################################################################
# MORE PARAMETERS (USUALLY NOT CHANGED)
```

```
# set paths to ImageJ and particle linker standalone
IJ.path <- "/home/felix/bin/ImageJ"
to.particlelinker <- "/home/felix/bin/ParticleLinker"

# directories and file names
to.data <- paste(getwd(),"/",sep="")
video.description.folder <- "0_video_description/"
video.description.file <- "video_description.txt"
raw.video.folder <- "1_raw/"
particle.data.folder <- "2_particle_data/"
trajectory.data.folder <- "3_trajectory_data/"
temp.overlay.folder <- "4a_temp_overlays/"
overlay.folder <- "4_overlays/"
merged.data.folder <- "5_merged_data/"
ijmacs.folder <- "ijmacs/"

# RAM allocation
memory.alloc <- c(60000)

# RAM per particle linker instance
memory.alloc.perLinker <- c(10000)

######################################################################
# VIDEO ANALYSIS

# identify particles
locate_and_measure_particles(to.data, raw.video.folder, particle.data.folder,
difference.lag, thresholds, min_size = particle_min_size, max_size =
```

```
particle_max_size, IJ.path, memory.alloc)

# link the particles
link_particles(to.data, particle.data.folder, trajectory.data.folder, linkrange =
trajectory_link_range, disp = trajectory_displacement, start_vid = 1, memory =
memory.alloc, memory_per_linkerProcess = memory.alloc.perLinker)

# merge info from description file and data
merge_data(to.data, particle.data.folder, trajectory.data.folder,
video.description.folder, video.description.file, merged.data.folder)

# load the merged data
load(paste0(to.data, merged.data.folder, "Master.RData"))

# filter data: minimum net displacement, their duration, the detection
#frequency and the median step length
trajectory.data.filtered <- filter_data(trajectory.data, filter_min_net_disp,
filter_min_duration, filter_detection_freq, filter_median_step_length)

# summarize trajectory data to individual-based data
morph_mvt <- summarize_trajectories(trajectory.data.filtered, calculate.median=F,
write = T, to.data, merged.data.folder)

# get sample level info
summarize_populations(trajectory.data.filtered, morph_mvt, write=T, to.data,
merged.data.folder, video.description.folder, video.description.file, total_frames)

# create overlays for validation
create_overlays(trajectory.data.filtered, to.data, merged.data.folder,
```

```
raw.video.folder, temp.overlay.folder, overlay.folder, 2048, 2048,
difference.lag, type = "label", predict_spec = F, IJ.path,
contrast.enhancement = 1, memory = memory.alloc)
```

### S3 Beverton-Holt model fitting

We ran population growth models with vaguely informative priors (i.e. mean estimates matched to observed data but with broad enough standard deviation to avoid constraining models). Model estimations were done using log-transformed parameters. Following parameters and priors were used for model fitting:

- $K.prior = log(1e5)$
- $K.sd.prior = 0.5$
- $r0.prior = -2.3$
- $r0.sd.prior = 0.5$
- $d.prior = -2.3$
- $d.sd.prior = 1.5$
- $N0.prior = log(1e3)$
- $N0.sd.prior = 0.5$
- $iter = 1e4$
- $warmup = 1e3$

### S4 Additional results

#### S4.1 Model comparison and summary table local adaptation test

Table S3: Model comparison table (of the dredge function) for the model for evolution of intrinsic rate of increase $r_0$ using AICc comparison.

| | **Intercept** | **Gene flow** | **Abiotic conditions** | **Reproduction** | **Gene flow×Abiotic conditions** | **Gene flow×Reproduction** | **Abiotic conditions×Reproduction** | **Gene flow×Abiotic conditions×Reproduction** | **df** | **logLik** | **AICc** | **delta** | **weight** |
|---|---|---|---|---|---|---|---|---|---|---|---|---|---|
| 24 | 0.0260 | + | + | + | | + | | | 6 | 18.0995 | -21.0878 | 0 | 0.32 |
| 32 | -0.0009 | + | + | + | + | + | | | 7 | 19.3827 | -20.4577 | 0.6300 | 0.24 |
| 56 | 0.009 | + | + | + | | + | + | | 7 | 18.7784 | -19.2492 | 1.8387 | 0.13 |
| 64 | -0.0223 | + | + | + | + | + | + | | 8 | 20.3339 | -18.9078 | 2.1800 | 0.11 |
| 3 | 0.1085 | | + | | | | | | 3 | 12.5652 | -18.3304 | 2.7573 | 0.08 |
| 7 | 0.1268 | | + | + | | | | | 4 | 12.7706 | -16.1620 | 4.9258 | 0.023 |
| 4 | 0.1064 | + | + | | | | | | 4 | 12.5679 | -15.7564 | 5.3314 | 0.021 |
| 128 | -0.0104 | + | + | + | + | + | + | + | 9 | 20.5450 | -15.5900 | 5.4978 | 0.02 |
| 39 | 0.0932 | | + | + | | | + | | 5 | 13.7973 | -15.4517 | 5.6361 | 0.02 |
| 12 | 0.0742 | + | + | | + | | | | 5 | 13.5422 | -14.9415 | 6.1463 | 0.01 |
| 8 | 0.1258 | + | + | + | | | | | 5 | 12.7713 | -13.3996 | 7.6882 | 0.01 |
| 16 | 0.0955 | + | + | + | + | | | | 6 | 13.8353 | -12.5595 | 8.5283 | 0.00 |
| 40 | 0.0951 | + | + | + | | | + | | 6 | 13.7996 | -12.4881 | 8.5997 | 0.00 |
| 48 | 0.0588 | + | + | + | + | | + | | 7 | 15.1617 | -12.0158 | 9.0720 | 0.00 |
| 1 | 0.3616 | | | | | | | | 2 | -12.1237 | 28.6346 | 49.7224 | 0.00 |
| 2 | 0.3111 | + | | | | | | | 3 | -11.8343 | 30.4687 | 51.5565 | 0.00 |
| 5 | 0.3799 | | | + | | | | | 3 | -12.0759 | 30.9517 | 52.0395 | 0.00 |

| | | | | | | | | | | | | | |
|---|---|---|---|---|---|---|---|---|---|---|---|---|---|
| 22 | 0.1978 | + | | + | | + | | | 5 | -10.016 | 32.1743 | 53.2620 | 0.00 |
| 6 | 0.3278 | + | | + | | | | | 4 | -11.7986 | 32.9766 | 54.0644 | 0.00 |

Table S4: Relative importance (RI) of the different independent variables obtained with AICc comparison of all possible models testing local adaptation as the evolution of intrinsic rate of increase $r_0$.

| **Independent factor** | **RI** |
|---|---|
| Abiotic conditions | 0.998 |
| Gene flow | 0.898 |
| Reproduction | 0.880 |
| Reproduction×Gene flow | 0.815 |
| Abiotic conditions×Gene flow | 0.387 |
| Reproduction×Abiotic conditions | 0.284 |
| Abiotic conditions×Reproduction×Gene flow | 0.021 |

Table S5: Summary table of the best model for evolution of intrinsic rate of increase $r_0$ according to AICc model comparison.

| | **Estimate** | **Std. Error** | **t value** | **Pr(>\|t\|)** |
|---|---|---|---|---|
| (Intercept) | 0.0260 | 0.0602 | 0.432 | 0.6688 |
| Reproduction “Sexual” | 0.1587 | 0.0798 | 1.9890 | 0.0562 |
| Gene flow “Present” | 0.1808 | 0.0767 | 2.356 | 0.0254 |
| Abiotic conditions “Gradient” | 0.6013 | 0.0543 | 11.072 | <0.0001 |
| Reproduction “Sexual”×Gene flow “Present” | -0.3488 | 0.1068 | -3.267 | 0.0028 |

#### S4.2 Expansion rate

In order to assess the effect of gradient, sex and gene flow on expansion rate (as in total number of patches expanded during the range expansion experiment), we fit a linear model (‘stats’-package) with total distance expanded as a function of pH gradient (gradient/no gradient), sex (sex/no sex) and gene flow (gene flow/no gene flow). We then used the dredge function

('MuMin'-package, version 1.43.6) using AICc comparison to select the best model.

Table S6: Model comparison table (of the dredge function) for the model for expansion rate using AICc comparison.

| | Intercept | Gene flow | Abiotic conditions | Reproduction | Gene flow×Abiotic conditions | Gene flow×Reproduction | Abiotic conditions×Reproduction | Gene flow×Abiotic conditions×Reproduction | df | logLik | AICc | delta | weight |
|---|---|---|---|---|---|---|---|---|---|---|---|---|---|
| 3 | 22.9500 | | + | | | | | | 3 | -43.4833 | 93.7942 | 0.0000 | 0.34 |
| 7 | 22.7374 | | + | + | | | | | 4 | -42.5460 | 94.5205 | 0.7263 | 0.24 |
| 39 | 22.9000 | | + | + | | | + | | 5 | -41.6489 | 95.5200 | 1.7257 | 0.15 |
| 4 | 22.9822 | + | + | | | | | | 4 | -43.4636 | 96.3558 | 2.5616 | 0.10 |
| 8 | 22.7659 | + | + | + | | | | | 5 | -42.5302 | 97.2827 | 3.4884 | 0.06 |
| 40 | 22.9230 | + | + | + | | | + | | 6 | -41.6376 | 98.5060 | 4.7118 | 0.03 |
| 12 | 22.9000 | + | + | | + | | | | 5 | -43.2297 | 98.6816 | 4.8874 | 0.03 |
| 16 | 22.6905 | + | + | + | + | | | | 6 | -42.3073 | 99.8453 | 6.0510 | 0.02 |
| 24 | 22.6933 | + | + | + | | + | | | 6 | -42.4443 | 100.1194 | 6.3252 | 0.01 |
| 56 | 22.8182 | + | + | + | | + | + | | 7 | -41.3965 | 101.2730 | 7.4787 | 0.01 |
| 48 | 22.8500 | + | + | + | + | | + | | 7 | -41.4312 | 101.3424 | 7.5482 | 0.01 |
| 32 | 22.6143 | + | + | + | + | + | | | 7 | -42.2138 | 102.9076 | 9.1134 | 0.00 |
| 64 | 22.7422 | + | + | + | + | + | + | | 8 | -41.1794 | 104.3587 | 10.5645 | 0.00 |
| 128 | 22.8000 | + | + | + | + | + | + | + | 9 | -41.0466 | 107.9194 | 14.1251 | 0.00 |
| 1 | 21.3939 | | | | | | | | 2 | -71.7947 | 147.9895 | 54.1953 | 0.00 |
| 2 | 21.8571 | + | | | | | | | 3 | -71.2102 | 149.2479 | 55.4537 | 0.00 |

| | | | | | | | | | | | | | |
|---|---|---|---|---|---|---|---|---|---|---|---|---|---|
| 5 | 21.1177 | | | + | | | | | 3 | -71.4974 | 149.8224 | 56.0282 | 0.00 |
| 6 | 21.5824 | + | | + | | | | | 4 | -70.9240 | 151.2766 | 57.4823 | 0.00 |
| 22 | 21.5714 | + | | + | | + | | | 5 | -70.9237 | 154.0695 | 60.2753 | 0.00 |

Table S7: Relative importance (RI) of the different independent variables obtained with AICc comparison of all possible models testing range expansion rate (total number of successful diserpsal events.

| **Independent factor** | **RI** |
|---|---|
| Gene flow | 0.273 |
| Abiotic conditions | 1 |
| Reproduction | 0.531 |
| Gene flow×Abiotic conditions | 0.061 |
| Gene flow×Reproduction | 0.029 |
| Abiotic conditions×Reproduction | 0.196 |
| Gene flow×Abiotic conditions×Reproduction | 0 |

Table S8: Type III ANOVA table of the best model for expansion rate, based on the AICc criterion.

| | **Degrees of freedom** | **F-value** | **Pr (>F)** |
|---|---|---|---|
| Abiotic conditions | 1 | 141.4 | <0.0001 |
| Residuals | 31 | | |

Table S9: Summary table of the best model for expansion rate, based on the AICc criterion.

| | **Estimate** | **Std. Error** | **t value** | **Pr(>\|t\|)** |
|---|---|---|---|---|
| (Intercept) | 22.950 | 0.209 | 110.08 | <0.0001 |
| Abiotic conditions "Gradient" | -3.950 | 0.332 | -11.89 | <0.0001 |

#### S4.3 Density dynamics during range expansion

We tested for changes in population density during the range expansion experiment by fitting a linear mixed model ('nlme'-package, version 3.1-137) of population density as a faction of 1) range expansion distance (number of successful dispersal events), 2) abiotic conditions, 3) reproduction and 4) gene flow, with replicate population as a random factor. We first fit a full interaction model using the maximum likelihood method, and then used the dredge function ('MuMin'-package, version 1.43.6) to find the best model based on the BIC (Bayesian Information Criterion) score. We then refit the best model using the restricted maximum likelihood method and used this best model to create model predictions. Due to the large size of the table, full model comparison is not included in this document, but model code to recreate the table is shown below. Only abiotic conditions, range expansion distance and their interaction strongly influenced population density (see table S10 for relative importances).

##### Model comparison code

```
full.model <- lme(data=data.all, dens.front~Gradient*Sex*Gene.Flow*pos,
random = ~1|ID, method = "ML")
comp <- dredge(full.model, rank = BIC)
best.model <- lme(data=data.all, dens.front~Gradient*pos,
random = ~1|ID, method = "REML")
```

Table S10: Relative importance (RI) of the different independent variables obtained with BIC comparison of all possible models testing population density during range expansion.

| **Independent factor** | **RI** |
|---|---|
| Gene flow | 0.266 |
| Abiotic conditions | 1 |
| Distance expanded (position) | 1 |
| Reproduction | 0.085 |
| Gene flow×Abiotic conditions | 0.011 |
| Gene flow×Range expansion distance | 0.012 |

| | |
|---|---|
| Gene flow×Reproduction | 0.001 |
| Abiotic conditions×Range expansion distance | 1 |
| Abiotic conditions×Reproduction | 0.003 |
| Position×Reproduction | 0.004 |
| Gene flow×Abiotic conditions×Range expansion distance | 0 |
| Gene flow×Abiotic conditions×Reproduction | 0 |
| Gene flow×Range expansion distance×Reproduction | 0 |
| Abiotic conditions×Range expansion distance×Reproduction | 0 |
| Gene flow×Abiotic conditions×Range expansion distance×Reproduction | 0 |

Table S11: Type III ANOVA table of the best model (fixed effects only) for population density dynamics, based on the BIC criterion.

| | **DF** | $\chi^2$ | **p-value** |
|---|---|---|---|
| (Intercept) | 1 | 660.541 | <0.0001 |
| Abiotic conditions | 1 | 0.044 | 0.833 |
| Range expansion distance | 1 | 4.526 | 0.034 |
| Abiotic conditions×Range expansion distance | 1 | 108.258 | <0.0001 |

Table S12: Summary table of the best model (fixed effects only) for population density dynamics, based on the BIC criterion.

| | **Value** | **Std.Error** | **DF** | **t-value** | **p-value** |
|---|---|---|---|---|---|
| (Intercept) | 165899.37 | 6454.979 | 746 | 25.701 | <0.0001 |
| Abiotic conditions "Gradient" | 2016.96 | 9581.401 | 38 | 0.211 0.834 | |
| Range expansion distance | -924.67 | 434.642 | 746 | -2.127 | 0.0337 |
| Abiotic conditions "Gradient"× Range expansion distance | -7244.62 | 696.283 | 746 | -10.405 | <0.0001 |

#### S4.4 Adaptation test raw evolution data

We tested differences using the raw intrinsic rate of increase $r_0$ (not standardized to the ancestor data by fitting a linear model ('stats'-package) with abiotic conditions ("Uniform" or "Gradient), reproduction ("Sexual", "Asexual") and gene flow ("Absent", "Present") as fixed factors. We fit the full interaction model, and then used the dredge function ('MuMin'-package, version 1.43.6) to compare all models using the AICc criterion.

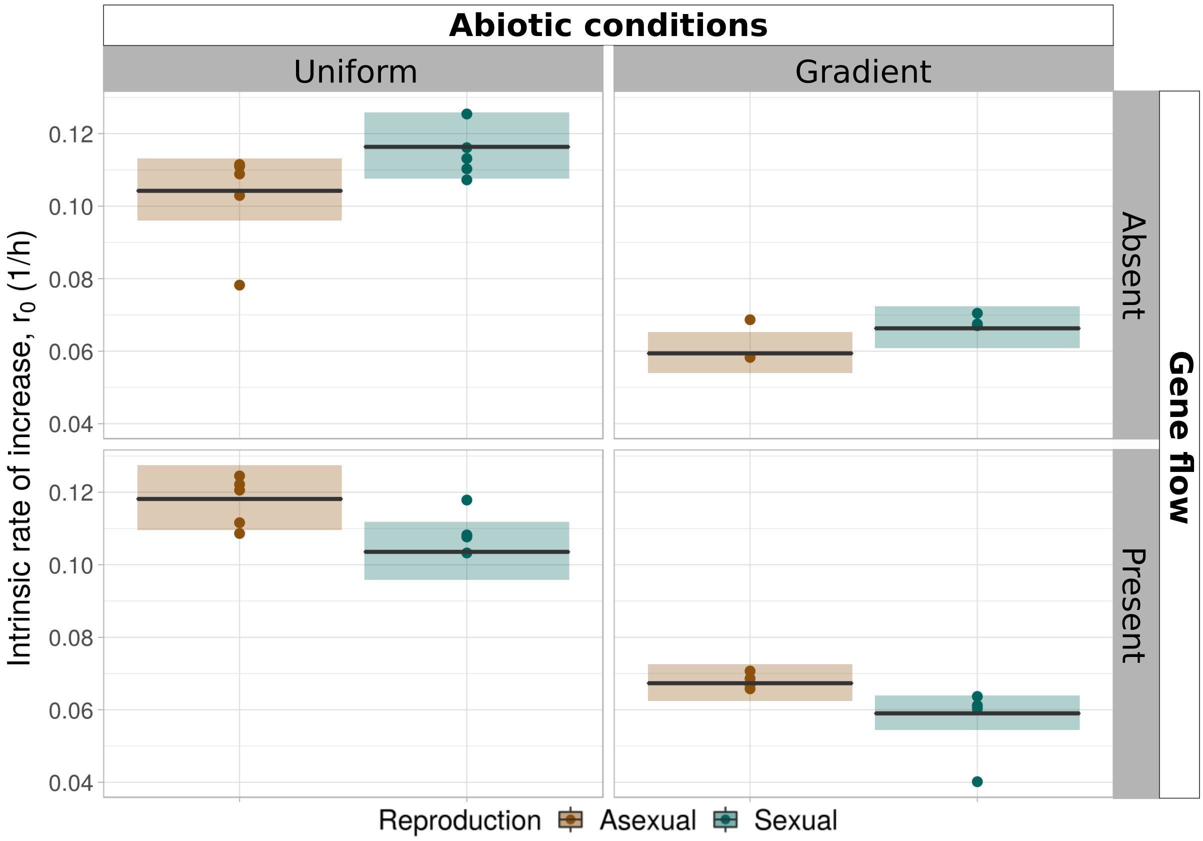


Figure S2: Evolution of intrinsic rate of increase $r_0$ based on the raw $r_0$-values. Y-axis shows $r_0$. Dots represent measurements, boxplots show the model predictions of the best model (mean and 95 % confidence intervals). Brown boxes and dots represent populations for treatment groups designated for asexual reproduction, green boxes and dots for treatment groups designated for sexual reproduction.

Table S13: Model comparison table (of the dredge function) for the model for evolution of intrinsic rate of increase $r_0$ based on the raw values, using AICc comparison.

| | Intercept | Gene flow | Abiotic conditions | Reproduction | Gene flow×Abiotic conditions | Gene flow×Reproduction | Abiotic conditions×Reproduction | Gene flow×Abiotic conditions×Reproduction | df | logLik | AICc | delta | weight |
|---|---|---|---|---|---|---|---|---|---|---|---|---|---|
| 24 | -2.2611 | + | + | + | | + | | | 6 | 30.5609 | -46.0107 | 0.0000 | 0.32 |
| 32 | -2.2798 | + | + | + | + | + | | | 7 | 31.8441 | -45.3806 | 0.6301 | 0.24 |
| 56 | -2.2728 | + | + | + | | + | + | | 7 | 31.2398 | -44.1720 | 1.8387 | 0.13 |
| 64 | -2.2946 | + | + | + | + | + | + | | 8 | 32.7953 | -43.8307 | 2.1800 | 0.11 |
| 3 | -2.2039 | | + | | | | | | 3 | 25.0266 | -43.2533 | 2.7574 | 0.08 |
| 7 | -2.1912 | | + | + | | | | | 4 | 25.2321 | -41.0848 | 4.9258 | 0.03 |
| 4 | -2.2054 | + | + | | | | | | 4 | 25.0293 | -40.6793 | 5.3314 | 0.02 |
| 128 | -2.2864 | + | + | + | + | + | + | + | 9 | 33.0065 | -40.5129 | 5.4978 | 0.02 |
| 39 | -2.2146 | | + | + | | | + | | 5 | 26.2587 | -40.3746 | 5.6361 | 0.02 |
| 12 | -2.2277 | + | + | | + | | | | 5 | 26.0036 | -39.8644 | 6.1463 | 0.01 |
| 8 | -2.1920 | + | + | + | | | | | 5 | 25.2327 | -38.3225 | 7.6882 | 0.01 |
| 16 | -2.2129 | + | + | + | + | | | | 6 | 26.2967 | -37.4824 | 8.5283 | 0.00 |
| 40 | -2.2132 | + | + | + | | | + | | 6 | 26.2611 | -37.4110 | 8.5997 | 0.00 |
| 48 | -2.2384 | + | + | + | + | | + | | 7 | 27.6232 | -36.9386 | 9.0720 | 0.00 |
| 1 | -2.4317 | | | | | | | | 2 | -6.8353 | 18.0578 | 64.0685 | 0.00 |
| 2 | -2.3899 | + | | | | | | | 3 | -6.5658 | 19.9316 | 65.9422 | 0.00 |
| 5 | -2.4190 | | | + | | | | | 3 | -6.8040 | 20.4079 | 66.4186 | 0.00 |
| 6 | -2.3740 | + | | + | | | | | 4 | -6.5218 | 22.4228 | 68.4335 | 0.00 |
| 22 | -2.4218 | + | | + | | + | | | 5 | -6.2065 | 24.5558 | 70.5665 | 0.00 |

Table S14: Type III ANOVA table of the best model for evolution of intrinsic rate of increase $r_0$ using raw $r_0$-values, based on the AICc criterion.

| | **Degrees of freedom** | **F-value** | **Pr (>F)** |
|---|---|---|---|
| Reproduction | 1 | 3.956 | 0.056 |
| Gene flow | 1 | 5.5525 | 0.025 |
| Abiotic conditions | 1 | 223.165 | <0.0001 |
| Reproduction×Gene flow | 1 | 10.675 | 0.003 |
| Residuals | 29 | | |

Table S15: Summary table of the best model for evolution of intrinsic rate of increase $r_0$ using raw $r_0$-values, based on the AICc criterion.

| | **Estimate** | **Std. Error** | **t value** | **Pr(>\|t\|)** |
|---|---|---|---|---|
| (Intercept) | -2.2611 | 0.0417 | -54.2000 | <.0001 |
| Reproduction “Sexual” | 0.1100 | 0.0553 | 1.9890 | 0.0562 |
| Gene flow “Present” | 0.1253 | 0.0532 | 2.356 | 0.0254 |
| Abiotic conditions “Gradient” | -0.5624 | 0.0376 | -14.9390 | <.0001 |
| Reproduction “Sexual”×Gene flow “Present” | -0.2418 | 0.0740 | -3.267 | 0.0028 |

#### S4.5 Evolution test over whole pH range

In order to test evolution of the entire pH-niche, we fit a linear mixed model (‘nlme’-packege, version 3.1-137) of change in intrinsic rate of increase $r_0$ as a function of 1) abiotic conditions (“Uniform”. “Gradient”), 2) reproduction (“Asexual”. “Sexual”), 3) gene flow (“Absent”. “Present”) and 4) pH of the assay medium, using population ID as a random effect. To account for potential non-linear responses to pH, we fit pH of the assay medium as a factor (pHfact). Change in intrinsic rate of increase was calculated as before, by dividing the $r_0$ of an assay culture by the mean value of the ancestors, and subsequently calculating the logarithm (base 2) of this ratio. We first fit the full interaction model based on the maximum likelihood method, and then used the dredge function (‘MuMin’-package, version 1.43.6) with BIC criterion to determine the best model. We then refit this model using the restricted maximum likelihood method to obtain model estimates and model predictions. Due to the table size, a full model comparison is not included, but the code used to obtain this table is listed below. Model comparison resulted in one clearly preferred model (model weight = 0.713; all other models had $\delta$BIC $>2$), with gradient and pHfact as the only included fixed effects. Note that only data with pH of 4.0 or higher is used, as ancestors typically only survived up to pH 4.0, and it was hence not possible for lower pH values to compare intrinsic rate of increase between ancestors and evolved populations.

**Model comparison code:**

full.model <- lme(data = dd.all, logratio ∼ Gradient*Sex*Gene.Flow*pHfact, random = ∼ 1|ID, method = ”ML”)

t.all <- dredge(full.model, rank = BIC)

best.model <- lme(data = dd.all, logratio ∼ Gradient + pHfact, random = ∼ 1|ID, method = ”REML”)

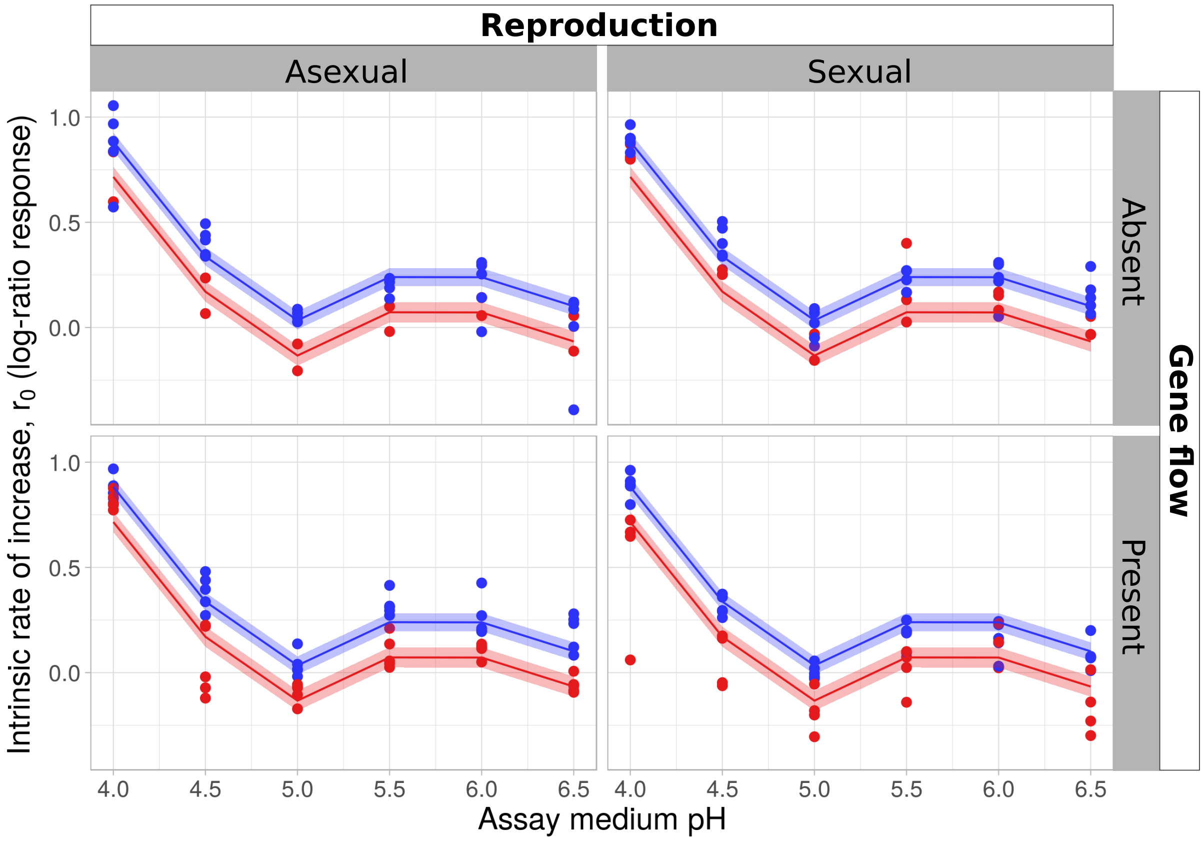


Figure S3: Evolution of intrinsic rate of increase $r_0$ over all pH values. The y-axis shows the change in $r_0$ compared to the ancestor, calculated as the logarithm (base 2) of the $r_0$ of the evolved population divided by the mean $r_0$ of the ancestors. Dots represent measurements, Lines and shaded areas show the model predictions of the best model (mean and 95 % confidence intervals). Blue lines, shaded areas and dots represent data and model predictions for the populations evolved without a pH gradient, red lines, shaded areas and dots for populations evolving with a pH gradient.

Table S16: Type III ANOVA table of the best model (fixed effects only) for evolution of intrinsic rate of increase $r_0$ over all measured pH values, based on the BIC criterion.

| | DF | F-value | p-value |
|---|---|---|---|
| (Intercept) | 1 | 356.2741 | <.0001 |
| Abiotic conditions | 1 | 43.063 | <.0001 |
| pH of assay medium (factorial) | 5 | 1796.068 | <.0001 |

Table S17: Summary table (fixed effects only) of the best model for evolution of intrinsic rate of increase $r_0$ over all measured pH values, based on the BIC criterion.

| | Value | Std.Error | DF | t-value | p-value |
|---|---|---|---|---|---|
| (Intercept) | 0.8828 | 0.0219 | 165 | 40.2775 | <.0001 |

| | | | | | |
|---|---|---|---|---|---|
| Abiotic conditions “Gradient” | -0.1673 | 0.0255 | 32 | -6.5622 | <.0001 |
| pH of assay medium (4.5) | -0.5452 | 0.02256 | 165 | -24.1297 | <.0001 |
| pH of assay medium (5) | -0.8486 | 0.0226 | 165 | -37.5602 | <.0001 |
| pH of assay medium (5.5) | -0.6431 | 0.0226 | 165 | -28.4639 | <.0001 |
| pH of assay medium (6) | -0.6438 | 0.0226 | 165 | -28.4958 | <.0001 |
| pH of assay medium (6.5) | -0.7816 | 0.0226 | 165 | -34.5944 | <.0001 |

#### S4.6 Survival probability

We measured survival by assessing at the end of the experimental range expansion, how many populations survived until the common garden stage, as populations sometimes died out after colonizing a new patch in the two-patch landscape, especially when expanding into a gradient. In order to test how survival of populations was affected by 1) abiotic conditions (“Uniform”/“Gradient”), 2) reproduction (“Asexual”/“Sexual”) and 3) gene flow (“Absent”/“Present”), we used generalized linear models with survival as a binomial response variable. Due to limitations in the data, where many one (1/survived) values were present and only few zero (0/extinct), we used Bayesian generalized linear models, as frequentist models did not converge properly when estimating the parameters, resulting in inflated standard deviation estimates. We created all possible models using the ‘Rethinking’-package (version 1.59), starting from a full interaction model to the intercept models, and then weighted the models using the WAIC criterion. We used the weighted estimates to calculate relative importance (RI) of the independent factors and to create weighted predictions, as show in figure S4. We used vaguely informative priors, meaning we included estimates that approximately corresponded to expected values, but including broad enough standard deviations, so models were not constrained too strongly in convergence. Below follows a list of the prior information and model structure:

- Survival probability $\sim$ binom(1, p)
- p = intercept + independent factors
- intercept $\sim$ dnorm(0, 1)

- All other independent factors $\sim$ dnorm(0, 10)

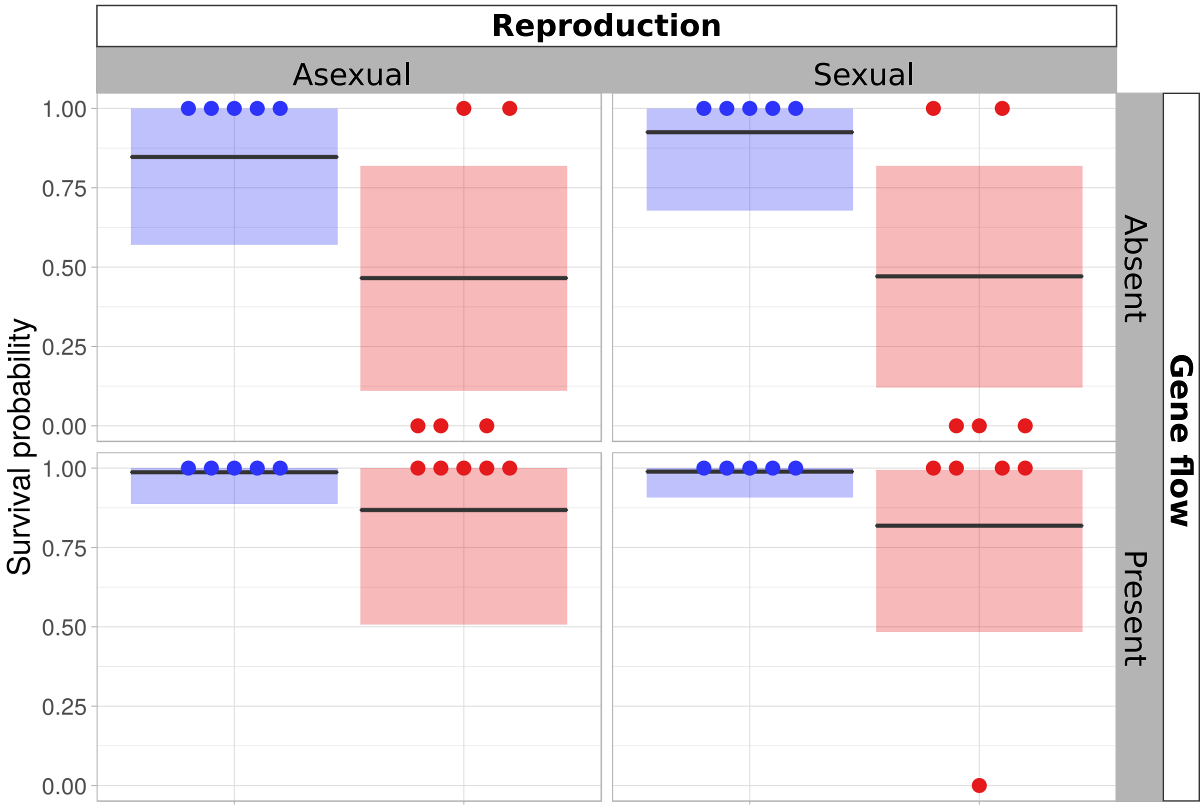


Figure S4: Survival of populations at the end of the range expansion experiment. Dots represent individual datapoints of populations that either survived (1) or went extinct (0). The boxes show the weighted predictions of survival based on the WAIC weighting on all Bayesian survival models, with the line representing the mean, and the range of the boxes the 95 % probability interval. Blue dots and boxes represent the populations expanding in a landscape without uniform abiotic conditions and red dots and boxes the populations expanding into a gradient.

Table S18: Relative importance (RI) of the effect of the independent factors on the survival probability of the populations during range expansion.

| **Independent factor** | **RI** |
|---|---|
| Abiotic conditions | 1 |
| Gene flow | 0.95 |
| Reproduction | 0.68 |
| Reproduction×Gene flow | 0.41 |
| Abiotic conditions×Gene flow | 0.34 |
| Reproduction×Abiotic conditions | 0.35 |

| | |
|---|---|
| Abiotic conditions×Reproduction×Gene flow | 0.09 |

### S4.7 Morphology: cell size

We assessed morphology (cell size and elongation) as well as cell movement (movement speed and turning angles) by fitting linear mixed models for these four response variables, using all the data of the bioassays (i.e. using measurements at every timepoint of the bioassays) as a function of 1) abiotic conditions, 2) reproduction, 3) gene flow, 4) pH of the assay medium and 5) population density (standardized as the proportion of population equilibrium density). We used this approach to test whether there was a difference between the eight treatment groups, while at the same time accounting for plastic effects associated with population density and pH. We used both replicate during experimental evolution and replicate population during bioassays as random effects. We compared models using the Dredge function in the MuMin package, however, as for the full model, the total number of levels of the explanatory variables exceeded the number the dredge function can handle (>30), we resorted to an extra step for model comparison.

We first created the full interaction model with all five explanatory variables, and calculated the BIC score. Next, we created all interaction models where one of the variables had been dropped from the full interaction model, and used the dredge function to compare all simplified models based on the BIC criterion. We then compared over all models which model had lowest BIC score.

Model comparison code:

```
#Do manual comparison
{
  full.model.size.evo <- lme(data = dd.evo,
   major_mean ~ percK*testpH*Gradient*Sex*Gene.Flow,
   random =  list(ID=~1, curveID=~1), method = "ML")

  t1 <-  dredge(lme(data = dd.evo,
```

```
   major_mean ~ testpH*Gradient*Sex*Gene.Flow,
  random =  list(ID=~1, curveID=~1), method = "ML"), rank = BIC)

  t2 <-  dredge(lme(data = dd.evo,
   major_mean ~ percK*Gradient*Sex*Gene.Flow,
   random =  list(ID=~1, curveID=~1), method = "ML"), rank = BIC)

  t3 <-  dredge(lme(data = dd.evo,
   major_mean ~ percK*testpH*Sex*Gene.Flow,
   random =  list(ID=~1, curveID=~1), method = "ML"), rank = BIC)

  t4 <-  dredge(lme(data = dd.evo,
   major_mean ~ percK*testpH*Gradient*Sex,
   random =  list(ID=~1, curveID=~1), method = "ML"), rank = BIC)

  t5 <-  dredge(lme(data = dd.evo,
   major_mean ~ percK*testpH*Gradient*Gene.Flow,
   random =  list(ID=~1, curveID=~1), method = "ML"), rank = BIC)

  BIC(full.model.size.evo)
  t1[1, "BIC"]
  t2[1, "BIC"]
  t3[1, "BIC"]
  t4[1, "BIC"]
  t5[1, "BIC"]
}

#best model
best.model.size.evo <- lme(data = dd.evo,
```

```
  major_mean ~ percK*testpH ,
  random =  list(ID=~1, curveID=~1))

 anova(best.model.size.evo)
 summary(best.model.size.evo)
 Anova(best.model.size.evo, contrasts=list(topic=contr.sum, sys=contr.sum),
  type=3)

 plot(best.model.size.evo)
}
```

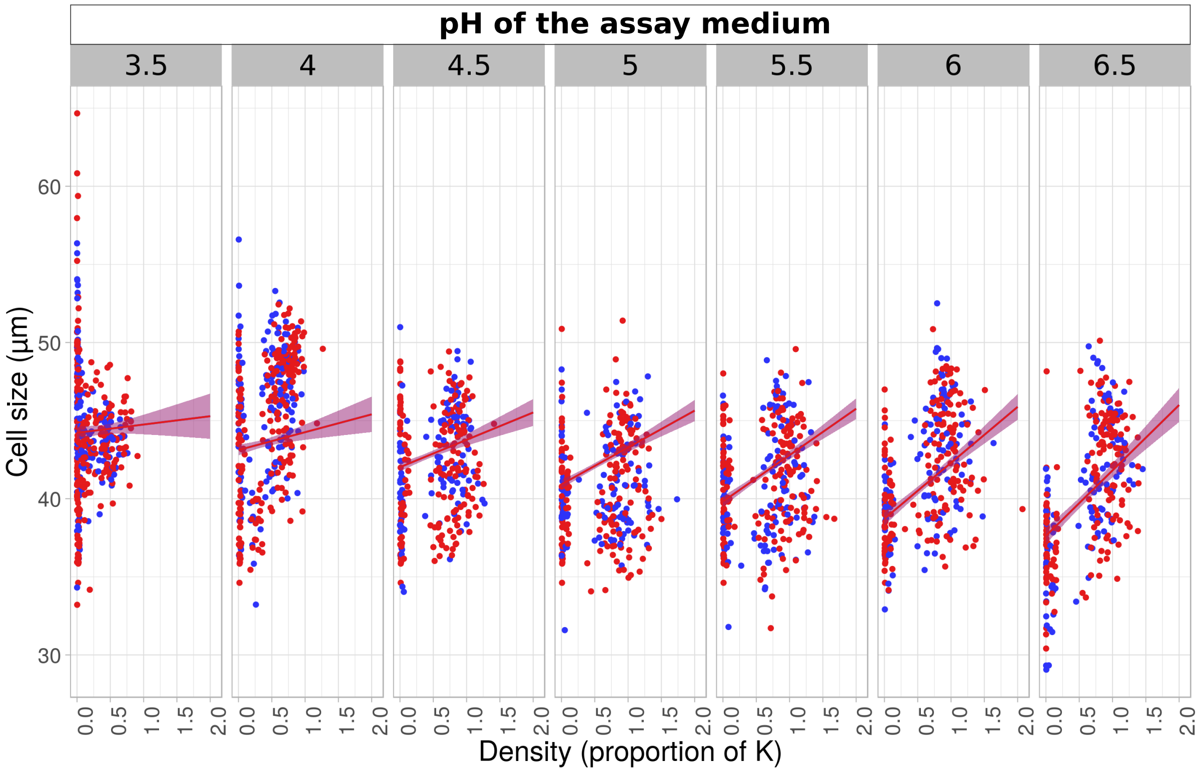


Figure S5: Cell size (longest axis through the cell) plotted to the standardized population density (percentage of the population equilibrium density K). Dots show datapoints, and full lines and shaded areas the model predictions (means and 95 % confidence intervals). Blue dots, lines and shaded areas represent data and predictions from populations that expanded into uniform abiotic conditions during the range expansion experiment. Red dots, lines and shaded areas show data and predictions for populations that expanded into a gradient during the range expansion experiment. The different panels represent the pH values of the assay medium in which the traits were measured.

Table S19: Type III ANOVA table of the best model for evolution of cell size, based on the BIC criterion.

| | F-value | Degrees of freedom | Pr (>F) |
|---|---|---|---|
| (Intercept) | 9.535 | 1 | 0.002 |
| pH of assay medium | 395.909 | 1 | <0.0001 |
| Density (prop. of K) | 4.941 | 1 | 0.026 |
| pH of assay medium×Density (prop. of K) | 24.277 | 1 | <0.0001 |

Table S20: Summary table of the best model for evolution of cell size, based on the BIC criterion.

| | Value | Std.Error | DF | t-value | p-value |
|---|---|---|---|---|---|
| (Intercept) | 51.911 | 0.662 | 2024 | 78.380 | <0.0001 |
| pH of assay medium | -2.198 | 0.136 | 200 | -16.220 | <0.0001 |
| Density (prop. of K) | -3.734 | 1.202 | 2024 | -3.105 | 0.002 |
| pH of assay medium×Density (prop. of K | 1.219 | 0.228 | 2024 | 5.33 | <0.0001 |

#### S4.8 Morphology: cell elongation

We assessed morphology (cell size and elongation) as well as cell movement (movement speed and turning angles) by fitting linear mixed models for these four response variables, using all the data of the bioassays (i.e. using measurements at every timepoint of the bioassays) as a function of 1) abiotic conditions, 2) reproduction, 3) gene flow, 4) pH of the assay medium and 5) population density (standardized as the proportion of population equilibrium density). We used this approach to test whether there was a difference between the eight treatment groups, while at the same time accounting for plastic effects associated with population density and pH. We used both replicate during experimental evolution and replicate population during bioassays as random effects. We compared models using the Dredge function in the MuMin package, however, as for the full model, the total number of levels of the explanatory variables exceeded the number the dredge function can handle (>30), we resorted to an extra step for model comparison.

We first created the full interaction model with all five explanatory variables, and calculated the BIC score. Next, we created all interaction models where one of the variables had been dropped from the full interaction model, and used the dredge function to compare all simplified models based on the BIC criterion. We then compared over all models which model had lowest BIC score.

Model comparison code:

```
#Do manual comparison
{
  full.model.elong.evo <- lme(data = dd.evo,
   major_mean/minor_mean ~ percK*testpH*Gradient*Sex*Gene.Flow,
   random =  list(ID=~1, curveID=~1), method = "ML")

  t1 <-  dredge(lme(data = dd.evo,
   major_mean/minor_mean ~ testpH*Gradient*Sex*Gene.Flow,
  random =  list(ID=~1, curveID=~1), method = "ML"), rank = BIC)

  t2 <-  dredge(lme(data = dd.evo,
   major_mean/minor_mean ~ percK*Gradient*Sex*Gene.Flow,
   random =  list(ID=~1, curveID=~1), method = "ML"), rank = BIC)

  t3 <-  dredge(lme(data = dd.evo,
   major_mean/minor_mean ~ percK*testpH*Sex*Gene.Flow,
   random =  list(ID=~1, curveID=~1), method = "ML"), rank = BIC)

  t4 <-  dredge(lme(data = dd.evo,
   major_mean/minor_mean ~ percK*testpH*Gradient*Sex,
   random =  list(ID=~1, curveID=~1), method = "ML"), rank = BIC)

  t5 <-  dredge(lme(data = dd.evo,
```

```
    major_mean/minor_mean ~ percK*testpH*Gradient*Gene.Flow,
    random =  list(ID=~1, curveID=~1), method = "ML"), rank = BIC)

   BIC(full.model.elong.evo)
   t1[1, "BIC"]
   t2[1, "BIC"]
   t3[1, "BIC"]
   t4[1, "BIC"]
   t5[1, "BIC"]
 }

 #best model
 best.model.elong.evo <- lme(data = dd.evo,
  major_mean/minor_mean ~ Gradient*percK + percK*testpH ,
  random =  list(ID=~1, curveID=~1))

 anova(best.model.elong.evo)
 summary(best.model.elong.evo)
 Anova(best.model.elong.evo, contrasts=list(topic=contr.sum, sys=contr.sum),
  type=3)

 plot(best.model.elong.evo)
}
```

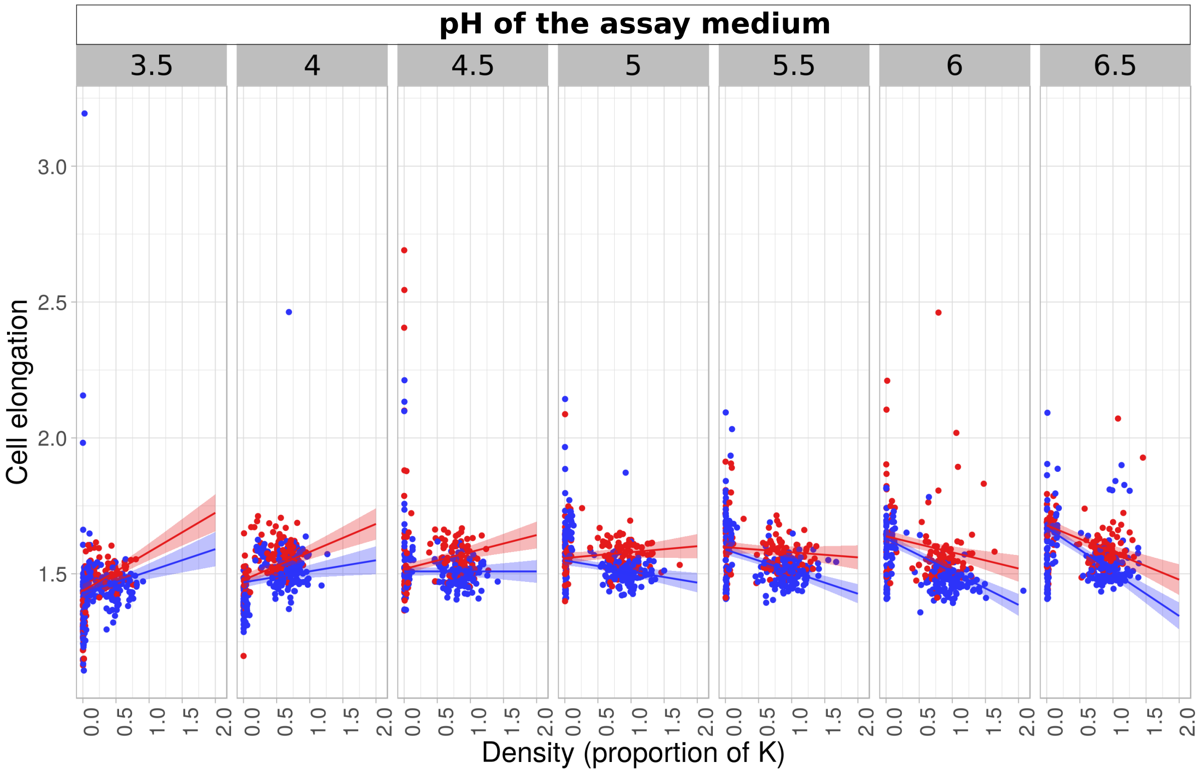


Figure S6: Cell elongation (ratio between longest and shortest axis through the cell) plotted to the standardized population density (percentage of the population equilibrium density K). Dots show datapoints, and full lines and shaded areas the model predictions (means and 95 % confidence intervals). Blue dots, lines and shaded areas represent data and predictions from populations that expanded into uniform abiotic conditions during the range expansion experiment. Red dots, lines and shaded areas show data and predictions for populations that expanded into a gradient during the range expansion experiment. The different panels represent the pH values of the assay medium in which the traits were measured.

Table S21: Type III ANOVA table of the best model for evolution of cell elongation, based on the BIC criterion.

| | **F-value** | **Degrees of freedom** | **Pr (>F)** |
|---|---|---|---|
| (Intercept) | 1692.8215 | 1 | <0.0001 |
| Abiotic conditions | 0.3924 | 1 | 0.5311 |
| Density (proportion of K) | 54.4074 | 1 | <0.0001 |
| pH of assay medium | 212.1209 | 1 | <0.0001 |
| Abiotic conditions “Gradient”×Density (prop. of K) | 12.6971 | 1 | 0.0004 |
| Density (proportion of K)×pH of assay medium | 76.2551 | 1 | <0.0001 |

Table S22: Summary table of the best model for evolution of cell elongation, based on the BIC criterion.

| | Value | Std.Error | DF | t-value | p-value |
|---|---|---|---|---|---|
| (Intercept) | 1.147 | 0.028 | 2023 | 41.143 | <0.0001 |
| Abiotic conditions "Gradient" | 0.009 | 0.014 | 32 | 0.626 | 0.5355 |
| Density (proportion of K) | 0.365 | 0.049 | 2023 | 7.376 | <0.0001 |
| pH of assay medium | 0.080 | 0.006 | 200 | 14.564 | <0.0001 |
| Abiotic conditions "Gradient"×Density (prop. of K) | 0.063 | 0.018 | 2023 | 3.563 | 0.0004 |
| Density (proportion of K)×pH of assay medium | -0.081 | 0.009 | 2023 | -8.732 | <0.0001 |

#### S4.9 Movement: cell speed

We assessed morphology (cell size and elongation) as well as cell movement (movement speed and turning angles) by fitting linear mixed models for these four response variables, using all the data of the bioassays (i.e. using measurements at every timepoint of the bioassays) as a function of 1) abiotic conditions, 2) reproduction, 3) gene flow, 4) pH of the assay medium and 5) population density (standardized as the proportion of population equilibrium density). We used this approach to test whether there was a difference between the eight treatment groups, while at the same time accounting for plastic effects associated with population density and pH. We used both replicate during experimental evolution and replicate population during bioassays as random effects. We compared models using the Dredge function in the MuMin package, however, as for the full model, the total number of levels of the explanatory variables exceeded the number the dredge function can handle (>30), we resorted to an extra step for model comparison.

We first created the full interaction model with all five explanatory variables, and calculated the BIC score. Next, we created all interaction models where one of the variables had been dropped from the full interaction model, and used the dredge function to compare all simplified models based on the BIC criterion. We then compared over all models which model had lowest BIC score.

Model comparison code:

```
#Do manual comparison
{
  full.model.speed.evo <- lme(data = dd.evo,
   gross_speed_mean ~ percK*testpH*Gradient*Sex*Gene.Flow,
   random =  list(ID=~1, curveID=~1), method = "ML")

  t1 <-  dredge(lme(data = dd.evo,
   gross_speed_mean ~ testpH*Gradient*Sex*Gene.Flow,
  random =  list(ID=~1, curveID=~1), method = "ML"), rank = BIC)

  t2 <-  dredge(lme(data = dd.evo,
   gross_speed_mean ~ percK*Gradient*Sex*Gene.Flow,
   random =  list(ID=~1, curveID=~1), method = "ML"), rank = BIC)

  t3 <-  dredge(lme(data = dd.evo,
   gross_speed_mean ~ percK*testpH*Sex*Gene.Flow,
   random =  list(ID=~1, curveID=~1), method = "ML"), rank = BIC)

  t4 <-  dredge(lme(data = dd.evo,
   gross_speed_mean ~ percK*testpH*Gradient*Sex,
   random =  list(ID=~1, curveID=~1), method = "ML"), rank = BIC)

  t5 <-  dredge(lme(data = dd.evo,
   gross_speed_mean ~ percK*testpH*Gradient*Gene.Flow,
   random =  list(ID=~1, curveID=~1), method = "ML"), rank = BIC)

  BIC(full.model.speed.evo)
  t1[1, "BIC"]
  t2[1, "BIC"]
```

```
    t3[1, "BIC"]
    t4[1, "BIC"]
    t5[1, "BIC"]
  }

  #best model
  best.model.speed.evo <- lme(data = dd.evo,
   gross_speed_mean ~ Gradient*percK + percK*testpH,
   random =  list(ID=~1, curveID=~1))

  anova(best.model.speed.evo)
  summary(best.model.speed.evo)
  Anova(best.model.speed.evo, contrasts=list(topic=contr.sum, sys=contr.sum),
   type=3)

  plot(best.model.speed.evo)
}
```

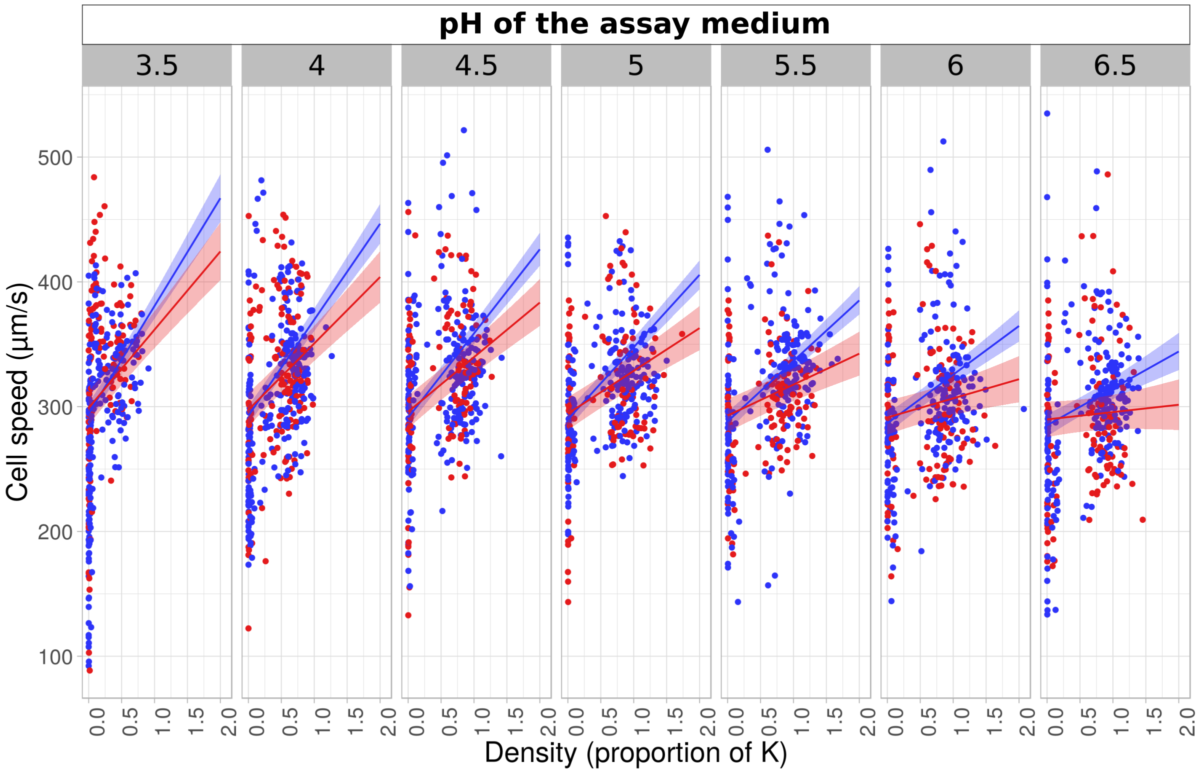


Figure S7: Cell swimming speed plotted to the standardized population density (percentage of the population equilibrium density K). Dots show datapoints, and full lines and shaded areas the model predictions (means and 95 % confidence intervals). Blue dots, lines and shaded areas represent data and predictions from populations that expanded into uniform abiotic conditions during the range expansion experiment. Red dots, lines and shaded areas show data and predictions for populations that expanded into a gradient during the range expansion experiment. The different panels represent the pH values of the assay medium in which the traits were measured.

Table S23: Type III ANOVA table of the best model for evolution of cell movement speed, based on the BIC criterion.

| | **F-value** | **Degrees of freedom** | **Pr (>F)** |
|---|---|---|---|
| (Intercept) | 1282.0050 | 1 | <0.0001 |
| Abiotic conditions | 0.3693 | 1 | 0.543 |
| Density (prop. of K) | 117.2553 | 1 | <0.0001 |
| pH of assay medium | 3.8034 | 1 | 0.051 |
| Abiotic conditions×Density (prop. of K) | 21.7046 | 1 | <0.0001 |
| Density (prop. of K)×pH of assay medium | 51.4084 | 1 | <0.0001 |

Table S24: Summary table of the best model for evolution of cell movement speed, based on the BIC criterion.

| | Value | Std.Error | DF | t-value | p-value |
|---|---|---|---|---|---|
| (Intercept) | 305.137 | 8.522 | 2326 | 35.8050 | <0.0001 |
| Abiotic conditions “Gradient” | 4.680 | 7.701 | 73 | 0.607 | 0.54 |
| Density (proportion of K) | 152.605 | 14.093 | 2326 | 10.829 | <0.0001 |
| pH of assay medium | -3.083 | 1.581 | 200 | -1.950 | 0.053 |
| Abiotic conditions “Gradient”×Density (prop. of K) | -23.676 | 5.082 | 2326 | -4.659 | <0.0001 |
| Density (proportion of K)×pH of assay medium | -18.932 | 2.640 | 2326 | -7.170 | <0.0001 |

#### S4.10 Movement: cell turning

We assessed morphology (cell size and elongation) as well as cell movement (movement speed and turning angles) by fitting linear mixed models for these four response variables, using all the data of the bioassays (i.e. using measurements at every timepoint of the bioassays) as a function of 1) abiotic conditions, 2) reproduction, 3) gene flow, 4) pH of the assay medium and 5) population density (standardized as the proportion of population equilibrium density). We used this approach to test whether there was a difference between the eight treatment groups, while at the same time accounting for plastic effects associated with population density and pH. We used both replicate during experimental evolution and replicate population during bioassays as random effects. We compared models using the Dredge function in the MuMin package, however, as for the full model, the total number of levels of the explanatory variables exceeded the number the dredge function can handle (>30), we resorted to an extra step for model comparison.

We first created the full interaction model with all five explanatory variables, and calculated the BIC score. Next, we created all interaction models where one of the variables had been dropped from the full interaction model, and used the dredge function to compare all simplified models based on the BIC criterion. We then compared over all models which model had lowest BIC score.

Model comparison code:

```
#Do manual comparison
```

```
{
  full.model.turning.evo <- lme(data = dd.evo,
   sd_turning_mean ~ percK*testpH*Gradient*Sex*Gene.Flow,
   random =  list(ID=~1, curveID=~1), method = "ML")

  t1 <-  dredge(lme(data = dd.evo,
   sd_turning_mean ~ testpH*Gradient*Sex*Gene.Flow,
  random =  list(ID=~1, curveID=~1), method = "ML"), rank = BIC)

  t2 <-  dredge(lme(data = dd.evo,
   sd_turning_mean ~ percK*Gradient*Sex*Gene.Flow,
   random =  list(ID=~1, curveID=~1), method = "ML"), rank = BIC)

  t3 <-  dredge(lme(data = dd.evo,
   sd_turning_mean ~ percK*testpH*Sex*Gene.Flow,
   random =  list(ID=~1, curveID=~1), method = "ML"), rank = BIC)

  t4 <-  dredge(lme(data = dd.evo,
   sd_turning_mean ~ percK*testpH*Gradient*Sex,
   random =  list(ID=~1, curveID=~1), method = "ML"), rank = BIC)

  t5 <-  dredge(lme(data = dd.evo,
   sd_turning_mean ~ percK*testpH*Gradient*Gene.Flow,
   random =  list(ID=~1, curveID=~1), method = "ML"), rank = BIC)

  BIC(full.model.turning.evo)
  t1[1, "BIC"]
  t2[1, "BIC"]
  t3[1, "BIC"]
```

```
    t4[1, "BIC"]
    t5[1, "BIC"]
  }

  #best model
  best.model.turning.evo <- lme(data = dd.evo,
   sd_turning_mean ~ percK*testpH ,
   random =  list(ID=~1, curveID=~1))

  anova(best.model.turning.evo)
  summary(best.model.turning.evo)
  Anova(best.model.turning.evo, contrasts=list(topic=contr.sum, sys=contr.sum),
   type=3)

  plot(best.model.turning.evo)
}
```

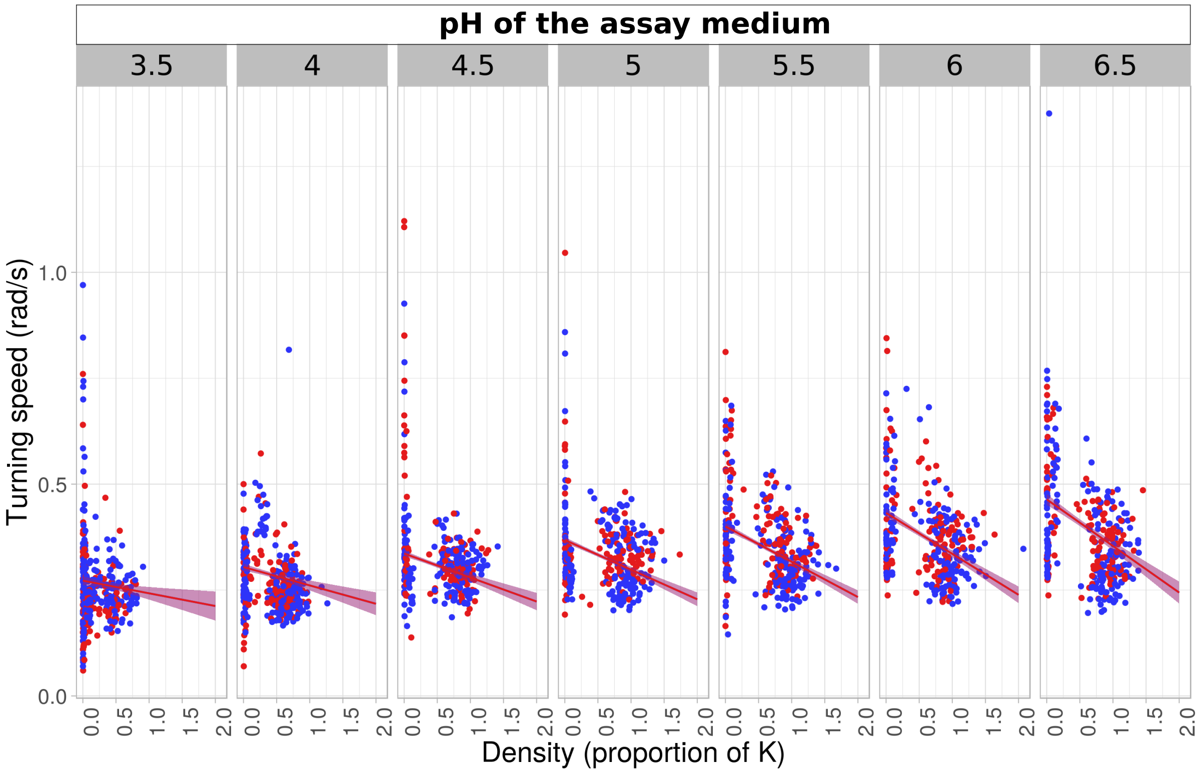


Figure S8: Cell turning speed plotted to the standardized population density (percentage of the population equilibrium density K). Dots show datapoints, and full lines and shaded areas the model predictions (means and 95 % confidence intervals). Blue dots, lines and shaded areas represent data and predictions from populations that expanded into uniform abiotic conditions during the range expansion experiment. Red dots, lines and shaded areas show data and predictions for populations that expanded into a gradient during the range expansion experiment. The different panels represent the pH values of the assay medium in which the traits were measured.

Table S25: Type III ANOVA table of the best model for evolution of cell turning speed, based on the BIC criterion.

| | **F-value** | **Degrees of freedom** | **Pr (>F)** |
|---|---|---|---|
| (Intercept) | 9.535 | 1 | 0.002 |
| pH of assay medium | 395.909 | 1 | <0.0001 |
| Density (prop. of K) | 4.941 | 1 | 0.026 |
| pH of assay medium×Density (prop. of K) | 24.277 | 1 | <0.0001 |

Table S26: Summary table of the best model for evolution of cell turning speed, based on the BIC criterion.

| | **Value** | **Std.Error** | **DF** | **t-value** | **p-value** |
|---|---|---|---|---|---|
| (Intercept) | 0.049 | 0.016 | 2024 | 3.088 | 0.002 |

| | | | | | |
|---|---|---|---|---|---|
| pH of assay medium | 0.064 | 0.003 | 200 | 19.897 | <0.0001 |
| Density (prop. of K) | 0.063 | 0.029 | 2024 | 2.223 | 0.026 |
| pH of assay medium×Density (prop. of K) | -0.027 | 0.005 | 2024 | -4.927 | <0.0001 |

#### S4.11 Dispersal rates

| | **Abiotic conditions** | **Reproduction** | **Gene flow** | **Mean percentage of dispersers** |
|---|---|---|---|---|
| **Treatment 1** | "Uniform" | "Asexual" | "Absent" | 6.00 |
| **Treatment 2** | "Uniform" | "Asexual" | "Present" | 4.98 |
| **Treatment 3** | "Uniform" | "Sexual" | "Absent" | 5.44 |
| **Treatment 4** | "Uniform" | "Sexual" | "Present" | 5.29 |
| **Treatment 5** | "Gradient" | "Asexual" | "Absent" | 3.22 |
| **Treatment 6** | "Gradient" | "Asexual" | "Present" | 2.49 |
| **Treatment 7** | "Gradient" | "Sexual" | "Absent" | 2.95 |
| **Treatment 8** | "Gradient" | "Sexual" | "Present" | 3.90 |

Table S27: Average dispersal rates (percentage of dispersers) for each of the eight treatment groups in the range expansion experiment.

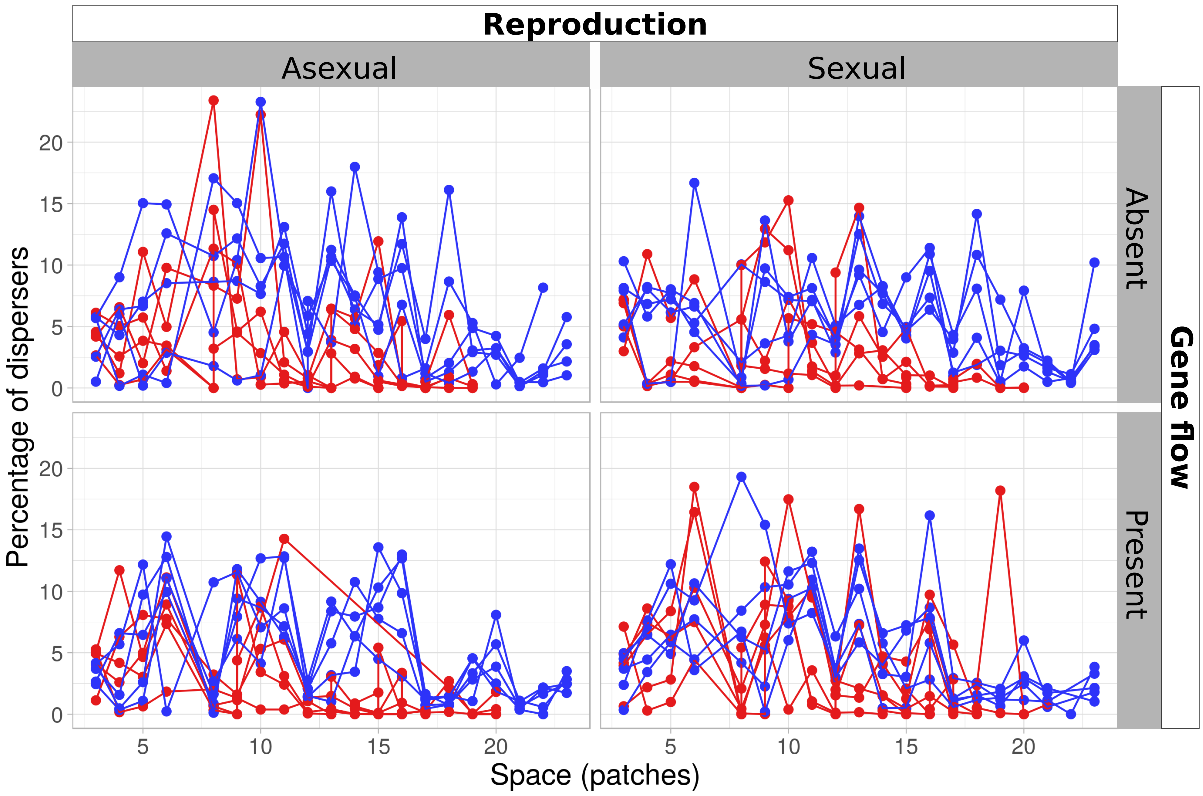


Figure S9: Dispersal rates (percentage of dispersers) plotted to the total distance expanded during range expansion. Blue dots and lines show data for populations expanding into uniform abiotic conditions, red dots and lines data for populations expanding into a gradient. Each line represents a distinct population over the course of the range expansion experiment.
